# Photoactivatable Rac1: a photosensor tilt by time-resolved crystallography and a partially open switch I when GDP-bound

**DOI:** 10.64898/2026.10.27.762334

**Authors:** Magdalena Pachota, Madan Kumar Shankar, Magdalena Wiśniewska, Anna Koczurowska, Cecilia Maria Casadei, Aleksandra Solecka, Sofia M. Kapetanaki, Bence Olasz, Larissa Lopes Cavalcante, Ran Ke, Dean Lang, Monika Bjelčić, Ayesha Asghar, Mirko Milas, Thomas Ursby, Manoop Chenchiliyan, Jie Nan, Edina Rosta, Przemyslaw Nogly

## Abstract

The LOV2 domain of *Avena sativa* phototropin 1 is widely used to control proteins with light, but how its signal reaches a fused effector has not been visualized. We followed photoactivatable Rac1 by time-resolved serial crystallography at room temperature. Within 350 µs, a fifth of the photosensors form the flavin–cysteine adduct and tilt as a rigid body on Rac1 about an axis through the domain interface, while the Jα helix stays docked and Rac1 keeps GTP and dark-state switches up to 1.85 ms. Dark crystals contain a variable GDP-bound sub-population, so light data must be referenced to interleaved dark frames. In this sub-population, switch I is only partially open, held at a Ca²⁺ ion that bridges LOV2 and Rac1, a conformation that can persist in simulations without the lattice. The photosensor thus first acts on Rac1 at the interface that also holds switch I in the GDP-bound state.

**Highlights:**

- Dark crystals vary in GDP content, so pump–probe data need interleaved dark frames
- The photosensor tilts on Rac1 within 350 µs about an axis through their interface
- The Jα helix stays docked and the Rac1 switches are unchanged up to 1.85 milliseconds
- After γ-phosphate loss, switch I is only partially open, held at the interface Ca²⁺

**eTOC blurb:** Pachota et al. follow the optogenetic tool photoactivatable Rac1 by time-resolved crystallography at room temperature. Within 350 µs, the photosensor tilts on its unchanged GTPase at their interface, and a GDP-bound sub-population in dark crystals shows switch I only partially open, held at a Ca²⁺ ion at the same interface.

## Introduction

Light-oxygen-voltage (LOV) domains sense blue light through a flavin mononucleotide (FMN), and light drives formation of a covalent adduct between the FMN C4a atom and a conserved cysteine within microseconds (Swartz *et al*., 2001; Konold *et al*., 2016). In the LOV2 domain of *Avena sativa* phototropin 1 (AsLOV2), the domain that primarily regulates phototropin kinase activity (Christie et al., 2002), adduct formation is coupled to unfolding of the C-terminal Jα helix. In the dark, Jα packs against the β-sheet of the PAS (Per-ARNT-Sim) fold (Harper *et al*., 2003), and the flanking Jα and A′α regions together modulate the light response (Halavaty & Moffat, 2007). Because the Jα helix provides a mechanical output that can be coupled to almost any partner, AsLOV2, a member of the widespread family of LOV photosensors (Glantz et al., 2016), has become a widely used building block of optogenetics, controlling enzyme activity and protein–protein interactions (Lee et al., 2008; Strickland et al., 2012; Wang et al., 2016), nuclear import (Niopek et al., 2014), organelle transport (van Bergeijk et al., 2015) and actin assembly by formins (Rao et al., 2013; Baarlink et al., 2013).

One of the first and most widely used of these tools is photoactivatable Rac1 (PA-Rac1; Wu *et al*., 2009). Rac1 is a Rho-family GTPase that regulates actin dynamics, cell protrusion and migration, largely through effectors such as the p21-activated kinases (Bosco *et al*., 2009; Bokoch, 2003). Like other small GTPases, it is active when bound to guanosine triphosphate (GTP) and inactive after hydrolysis of GTP to guanosine diphosphate (GDP), and it signals through two flexible regions, switch I and switch II, which bind effectors in the GTP-bound state and change conformation when the γ-phosphate is lost (Vetter & Wittinghofer, 2001). PA-Rac1 is a fusion protein in which AsLOV2 with its Jα helix (residues 404–546; LOV2 in the following) is attached to the N-terminus of Rac1 (Figure 1A), so that the LOV2 domain sterically blocks the effector-binding surface of Rac1 until light unwinds Jα. Rac1 in PA-Rac1 also carries the Q61L substitution, which abolishes GTP hydrolysis, and the E91H and N92H substitutions, which diminish its interactions with regulatory proteins (Wu *et al*., 2009). Cryogenic crystal structures of the dark state showed this arrangement (PDB entries 2WKP, 2WKQ and 2WKR; Wu et al., 2009), and the switch regions of Rac1 are known to be dynamic and sensitive to mutation (Acuner *et al*., 2021). In solution, however, illumination does not simply release Rac1 from the photosensor but shifts the population between pre-existing caged conformations, in which LOV2 blocks the effector-binding surface, and uncaged ones, and a Ca²⁺ ion bound at the LOV2–Rac1 interface stabilizes the caged Jα conformation in the dark (Winkler *et al*., 2015; Lindner *et al*., 2015).

**Figure 1.**
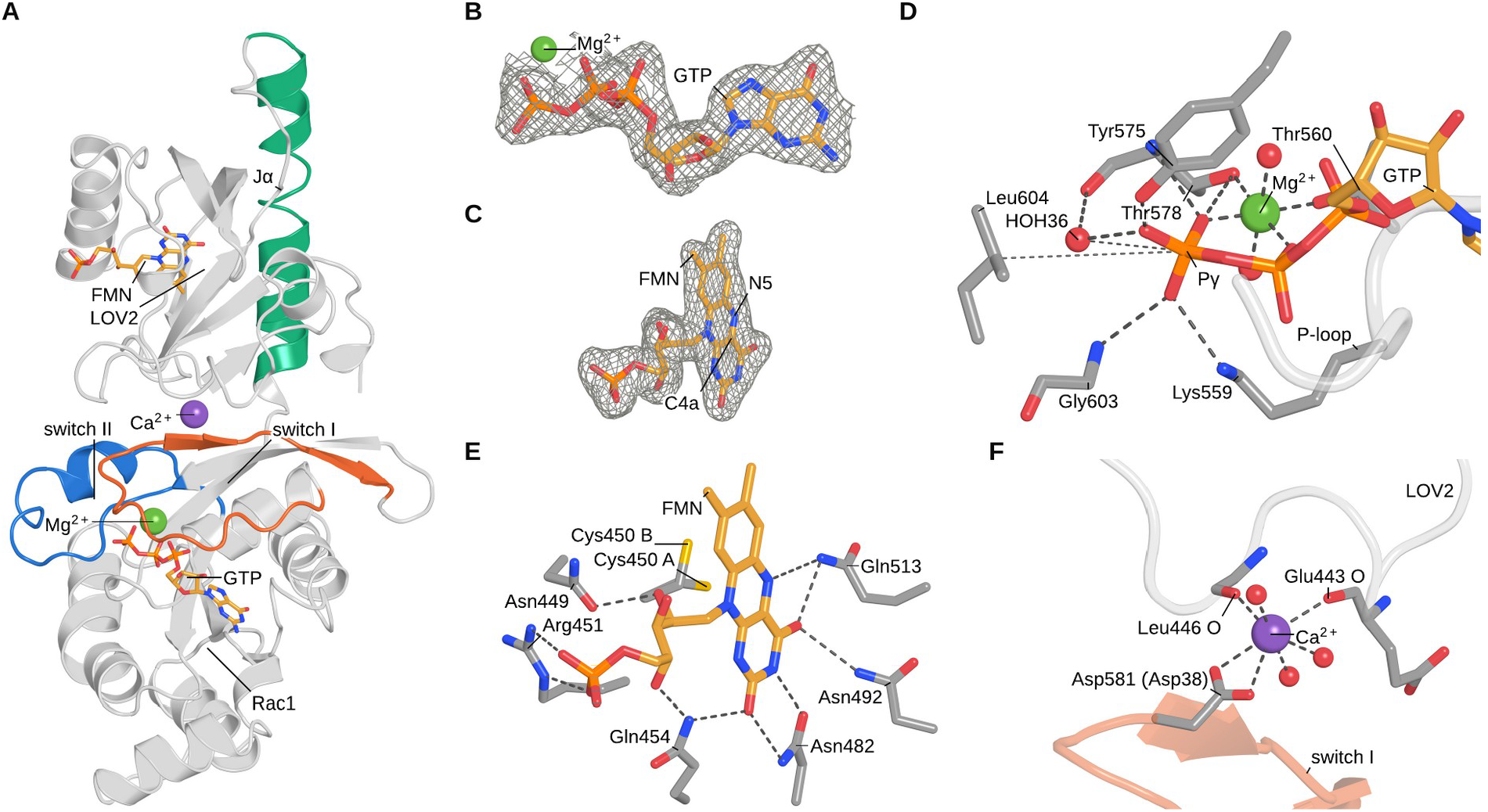
Structure of PA-Rac1 in the dark state (2026 interleaved dark, 2.01 Å). (A) Overall structure as a gray cartoon, with the Jα helix in green and Rac1 switch I (residues 569–588, Rac1 26–45) in orange and switch II (residues 600–619, Rac1 57–76) in blue; FMN and GTP as sticks and the Mg²⁺ and Ca²⁺ ions as spheres. (B) 2mFo–DFc density for GTP and Mg²⁺ and (C) for FMN at 1.0σ (0.18 e Å⁻s; σA-weighted map calculated with Servalcat). (D) γ-Phosphate and Mg²⁺ site: dashes, Mg²⁺ coordination (Thr560 OG1 2.12 Å, Thr578 OG1 2.18 Å, β- and γ-phosphate oxygens 2.01 and 1.91 Å, two water molecules 2.09 Å) and polar contacts of the γ-phosphate (Thr578 OG1 2.94 Å, Thr578 N 3.00 Å, Tyr575 OH 2.65 Å, Gly603 N 2.96 Å, Lys559 NZ 2.76 Å); dotted lines, from the γ-phosphorus to HOH36, the water closest to it (3.49 Å, close to in line with the bond to the bridging β-phosphate oxygen; O3B–PG–O 155°), and to Leu604 CG (the Q61L substitution; 5.9 Å). Guanine, ribose and P-loop contacts of the 2025 dark model are listed in Table S1. (E) Hydrogen-bond network of the FMN chromophore (Gln513 NE2–O4 3.14 Å), with Cys450 in both modeled rotamers (A, B; occupancies 0.45 and 0.55). (F) The interface Ca²⁺ ion and its seven ligands: the Glu443 and Leu446 carbonyl oxygens of LOV2, both carboxylate oxygens of Asp581 (Rac1 Asp38) in switch I and three water molecules (2.62, 2.40, 2.48, 2.81 and 2.23–2.50 Å). Carbon atoms of FMN and GTP are orange, those of the protein gray.

How the light signal travels from the flavin to Jα has been studied extensively by spectroscopy and solution scattering. Time-resolved infrared spectroscopy shows structural changes first in the β-sheet and then in the helical regions of AsLOV2 (Gil *et al*., 2017) and that Jα unfolds within about 500 µs (Konold *et al*., 2016). Time-resolved X-ray solution scattering has resolved an intermediate that precedes Jα unfolding (Konold *et al*., 2024) and shown that Jα unfolds with a time constant of about 0.7 ms (Kim et al., 2024). The conserved Gln513 has been proposed to couple protonation of the flavin N5 to the β-sheet (Nash *et al*., 2008; Iuliano *et al*., 2020), although LOV receptors lacking this glutamine can still signal (Dietler *et al*., 2022), and simulations suggest further pathways through the β-sheet (Peter *et al*., 2010, 2012; Arshi *et al*., 2023). In an engineered fusion of LOV2 with the GTPase Cdc42, NMR has shown that the light state perturbs the effector-binding site of the GTPase (Jain *et al*., 2023). These methods report on secondary structure, overall shape or individual sites rather than on the complete atomic structure, and whether and how the signal reaches a fused effector has not been observed directly by crystallography.

Time-resolved serial crystallography captures light-induced structural changes in protein microcrystals at room temperature (Brändén & Neutze, 2021). At synchrotrons, time-resolved serial synchrotron crystallography (TR-SSX) has captured millisecond intermediates of light-driven proteins (Weinert *et al*., 2019) and adduct formation and C-terminal changes of the isolated CrPhotLOV1 domain between 2.5 and 95 ms after excitation (Gotthard *et al*., 2024b), whereas serial oscillation crystallography has followed the build-up of the light state of Arabidopsis phototropin 2 LOV2 at 63 ms resolution (Aumonier et al., 2020). At X-ray free-electron lasers (XFELs), the light-induced structural changes of CrPhotLOV1 have been resolved 10 µs after excitation (Gotthard *et al*., 2024a) and from picoseconds to 100 µs (Olasz *et al*., 2026a). These studies used isolated LOV domains, without a Jα helix or a fused effector.

The MicroMAX beamline at MAX IV extends TR-SSX into the microsecond range (Gonzalez et al., 2025), allowing the early structural events of LOV2 activation to be resolved. Here we report TR-SSX of PA-Rac1 at MicroMAX to determine where the light signal first passes from the photosensor to the effector. We performed two experiments: in 2025, with nanosecond excitation and one merged light dataset, and in 2026, with a time series of four windows up to 1.85 ms after excitation. Each light dataset was referenced to the dark frames interleaved with the pump–probe frames of the same runs (interleaved dark); dark data recorded separately with the laser off (laser-off dark) were used to characterize the dark crystals. These structures extend time-resolved crystallography of LOV domains to a LOV2 photosensor with its Jα helix and a fused effector on the sub-millisecond timescale. Up to 1.85 ms after excitation, the light response reaches Rac1 only as a tilt of the photosensor about the domain interface. A GDP-bound sub-population in the dark crystals shows that switch I stays anchored at the same interface, at a Ca²⁺ ion that bridges LOV2 and Rac1, when the γ-phosphate is lost.

## Results

### Room-temperature structure of PA-Rac1 in the dark state

The room-temperature dark-state structure of PA-Rac1 was determined from the interleaved dark data of the 2026 experiment, which serve as the reference for its pump–probe data, and refined at 2.01 Å (Table 1). Residues 404–723 (LOV2 residues 404–546 and Rac1 residues 4–180; Rac1 numbering, given in parentheses throughout, is the fusion numbering minus 543) are resolved without chain breaks, and thirteen side chains, including that of the photoreactive Cys450, are modeled in two conformations. As in the cryogenic structures of PA-Rac1 (PDB entries 2WKP, 2WKQ and 2WKR; Wu *et al*., 2009), the Jα helix (Asp522–Ala543) packs against the β-sheet of the LOV2 domain, which shields the effector-binding surface of Rac1 (**Figure 1A**; secondary-structure assignment in Methods). The 2025 laser-off dark structure (2.43 Å), the reference for the 2025 data, superposes on it with a Cα root-mean-square deviation (RMSD) of 0.13 Å.

**Table 1.**
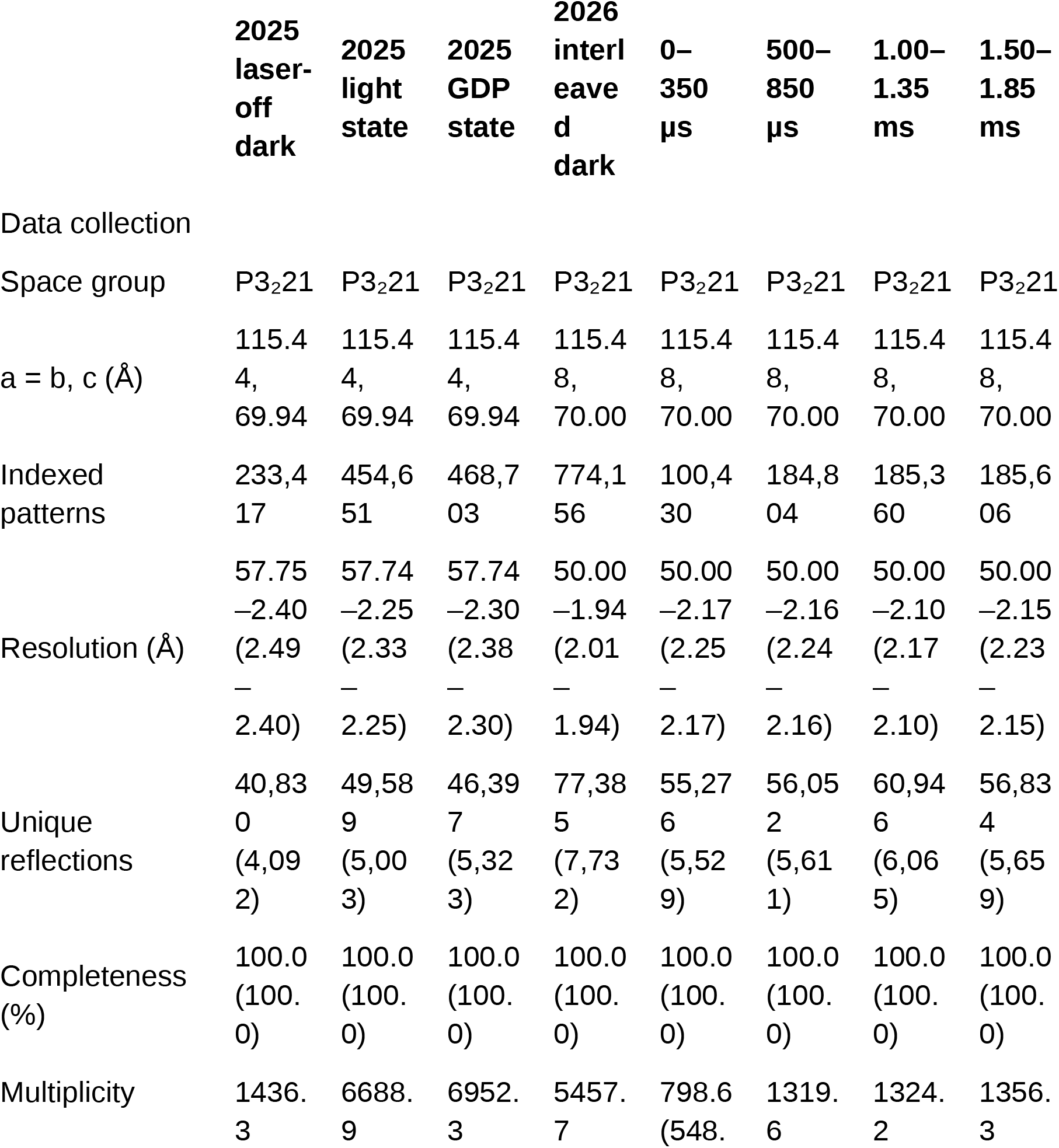
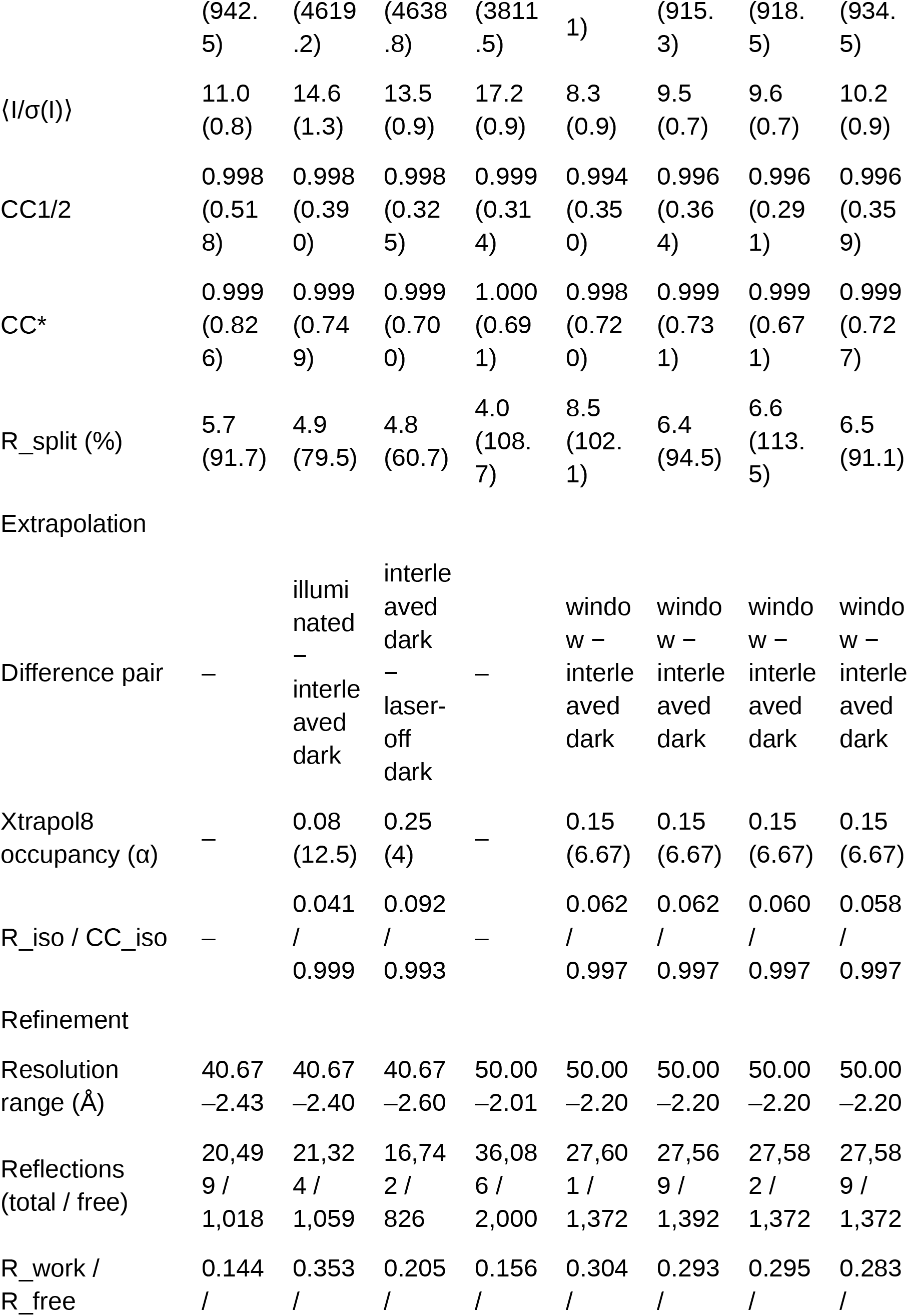
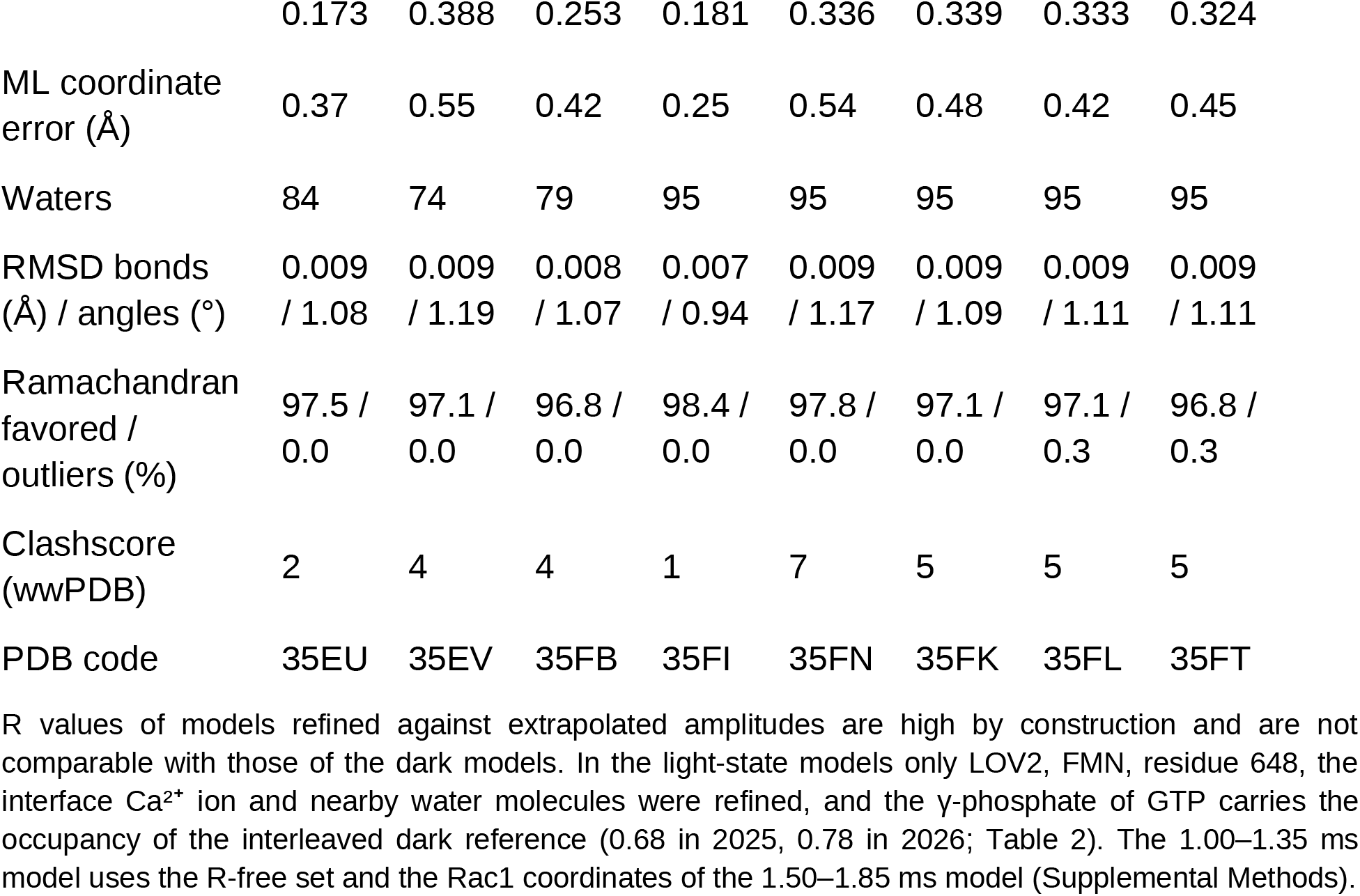
Data collection, extrapolation and refinement statistics. Data-collection statistics are from CrystFEL for the merged data of each dataset (for extrapolated entries, the triggered dataset) to the resolution given; values for the highest-resolution shell are in parentheses. Unique reflections and multiplicity count Friedel mates separately. CC* is defined as in Karplus & Diederichs (2012). For extrapolated entries, the number in parentheses after α is the extrapolation factor 1/α. Refinement statistics are from the final refinements in Phenix 1.21.2, and clashscores from the wwPDB validation reports.

FMN is held in the PAS core by a network of hydrogen bonds that includes the Gln513 amide, proposed to relay the signal to the β-sheet (Nash *et al*., 2008; Iuliano *et al*., 2020), and Cys450 occupies two rotamers (Figure 1C, E). Rac1 binds GTP and a Mg²⁺ ion with every canonical contact to the γ-phosphate of a GTP-bound small GTPase in place (Vetter & Wittinghofer, 2001), from Thr578 and Tyr575 (Rac1 Thr35 and Tyr32) in switch I and Gly603 (Rac1 Gly60) in switch II to Lys559 (Rac1 Lys16) in the P-loop (Figure 1B, D). No side chain takes the place of the catalytic Gln61, which the Q61L substitution removes from switch II.

The photosensor docks onto the effector-binding surface of Rac1: excluding the covalent junction (residues 544–549), sixteen LOV2 residues and fourteen Rac1 residues lie within 4 Å of the other domain, nine of them in switch I or switch II. At this interface, a Ca²⁺ ion bridges the two domains (Figure 1F). It is coordinated by the main-chain carbonyls of Glu443 and Leu446 in LOV2, by both carboxylate oxygens of Asp581 (Rac1 Asp38) at the C-terminal end of switch I and by three water molecules, 13.7 Å from the γ-phosphorus. The site is occupied in the dark datasets of both experiments and in the cryogenic structure 2WKP, whose crystals were grown, like ours, in a condition containing calcium acetate, whereas in crystals grown without calcium acetate a water molecule occupies this position (2WKQ and 2WKR; Wu et al., 2009).

### Matched dark references reveal a GDP-bound sub-population independent of illumination

Because the difference maps, calculated with Xtrapol8 (De Zitter *et al*., 2022), depend on the dark reference, we refined the dark datasets of both experiments with the same protocol (Table 2; Methods). None of them shows density for the photoadduct (at most 1.4σ in the 2mFo–DFc map at the position of the covalent Cys450 Sγ atom, against 6–11σ on the nucleotide phosphates), and the 2026 interleaved dark carries no detectable activation (implied activated fraction −0.006 ± 0.006 relative to the laser-off dark), so the light-minus-dark maps of the time series are not attenuated by stray light.

**Table 2.**
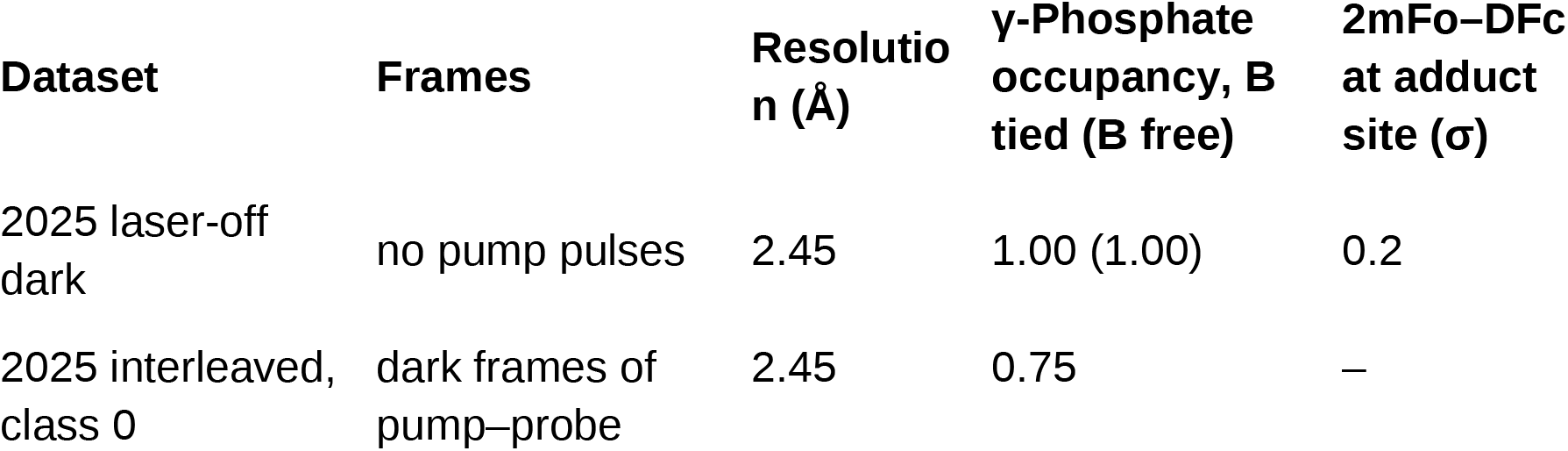
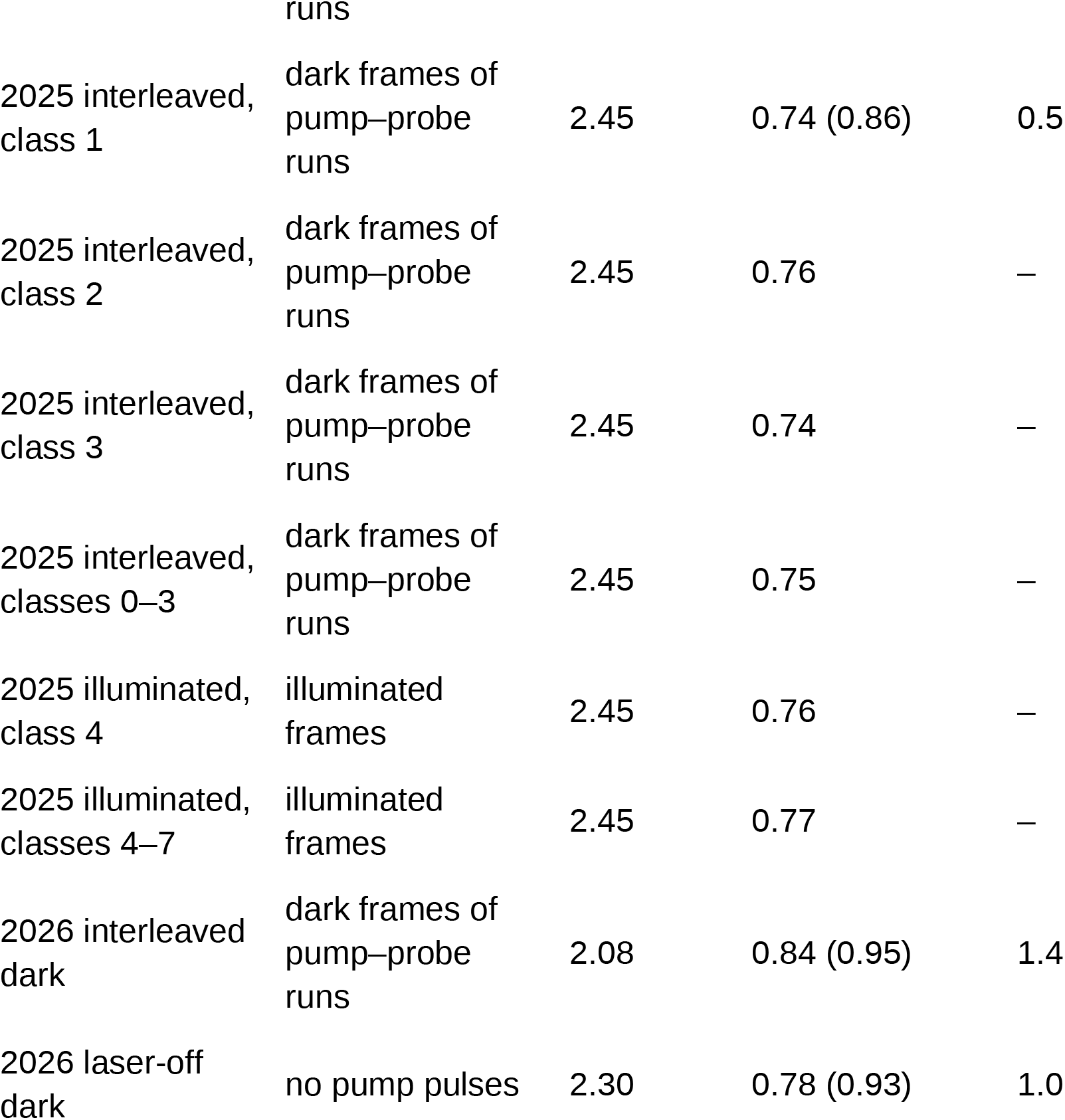
γ-Phosphate occupancy of GTP in the dark and light datasets refined with one protocol (Servalcat; Methods): 2025 datasets at 2.45 Å from the 2025 laser-off model, 2026 datasets at 2.08 Å (interleaved) and 2.30 Å (laser-off) from the 2026 dark model. The occupancy was refined with coordinates fixed and the B factor tied to that of the β-phosphorus and, in parentheses where available, with B factor and coordinates refined freely. The last column gives 2mFo–DFc at the position of the covalent Cys450 Sγ atom of the light-state model (adduct site), in σ units of the unit cell. Refined in Phenix as one occupancy group, the γ-phosphate occupancy is 0.98 (2025 laser-off dark), 0.68 (2025 interleaved dark), 0.78 (2026 interleaved dark) and 0.74 (2026 laser-off dark); the deposited 2026 dark model and the light-state models carry the value of the interleaved dark of the same experiment.

| Dataset | Frames | Resolution (Å) | γ-Phosphate occupancy, B tied (B free) | 2mFo–DFc at adduct site (σ) |
| --- | --- | --- | --- | --- |
| 2025 laser-off dark | no pump pulses | 2.45 | 1.00 (1.00) | 0.2 |
| 2025 interleaved, class 0 | dark frames of pump–probe runs | 2.45 | 0.75 | – |
| 2025 interleaved, class 1 | dark frames of pump–probe runs | 2.45 | 0.74 (0.86) | 0.5 |
| 2025 interleaved, class 2 | dark frames of pump–probe runs | 2.45 | 0.76 | – |
| 2025 interleaved, class 3 | dark frames of pump–probe runs | 2.45 | 0.74 | – |
| 2025 interleaved, classes 0–3 | dark frames of pump–probe runs | 2.45 | 0.75 | – |
| 2025 illuminated, class 4 | illuminated frames | 2.45 | 0.76 | – |
| 2025 illuminated, classes 4–7 | illuminated frames | 2.45 | 0.77 | – |
| 2026 interleaved dark | dark frames of pump–probe runs | 2.08 | 0.84 (0.95) | 1.4 |
| 2026 laser-off dark | no pump pulses | 2.30 | 0.78 (0.93) | 1.0 |

The dark datasets differ, however, at the nucleotide (Table 2; Figure 2A). With its B factor tied to that of the β-phosphorus, the γ-phosphate occupancy refines to 1.00 in the 2025 laser-off dark, but to 0.74–0.76 in each of the four interleaved dark classes of the same experiment, 0.84 in the 2026 interleaved dark and 0.78 in the 2026 laser-off dark. The crystals therefore contain a GDP-bound sub-population of up to about a quarter of the molecules, whose size differs between samples. The protein was purified and crystallized with GTP, and Q61L Rac1 does not hydrolyze GTP measurably (Davis et al., 2013), so how this sub-population forms is open (see Discussion). In the 2025 pump–probe runs it is the same in the dark frames (0.74–0.76) and the illuminated frames (0.76–0.77), so it reflects the history of the sample, not the pump pulse. The modification at Cys648 (Rac1 Cys105), modeled as S-sulfocysteine, follows the same pattern: it is weakest in the 2025 laser-off dark and gives features of +6 to +20σ in the dark-minus-dark maps. Because light and dark frames of each pump–probe experiment come from the same sample stream, the GDP-bound fraction cancels in the light-minus-dark maps.

**Figure 2.**
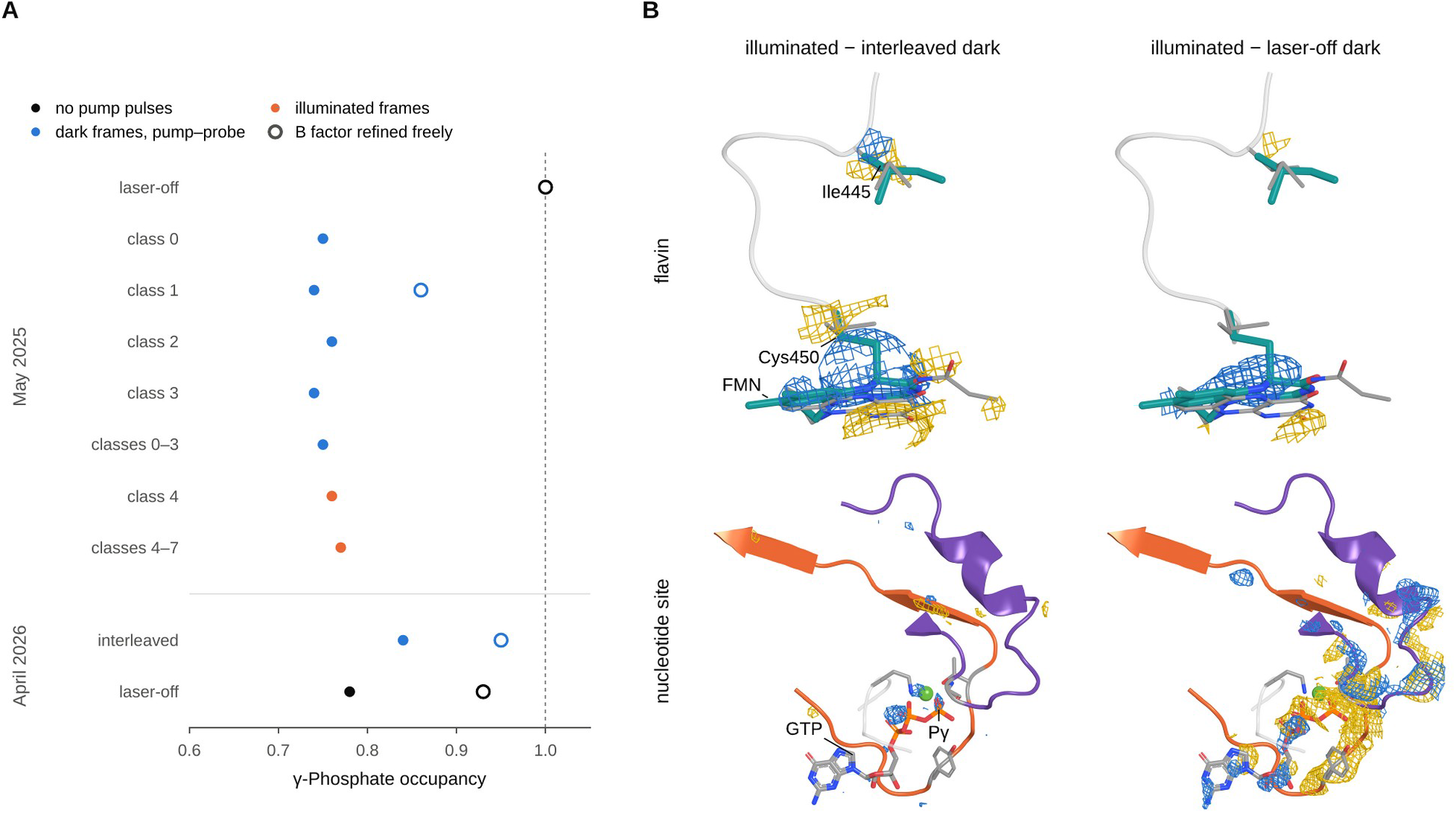
Dark references and the GDP-bound sub-population. (A) γ-Phosphate occupancy refined with the same protocol for the dark and light datasets (Table 2): datasets recorded without pump pulses (black), dark frames of pump–probe runs (blue) and illuminated frames (orange); filled symbols, B factor tied to that of the β-phosphorus; open symbols, B factor refined freely. (B) The 2025 light data referenced to the interleaved dark of the same runs (illuminated minus interleaved dark, left) and to the separately recorded laser-off dark (illuminated minus laser-off dark, right), at the flavin (top) and at the Rac1 nucleotide site (bottom). Xtrapol8 difference maps contoured at ±3.0σ; positive density in blue, negative in gold; models and views as in Figures 3E and 5B.

Referencing to a separately recorded dark would instead have misrepresented the light response. In 2025, the illuminated minus laser-off map is dominated by the GDP-bound sub-population (−20.0σ at the γ-phosphorus) and shows only weak density at the photoadduct (+1.7σ at the adduct Sγ position, +4.8σ within 1 Å), whereas the illuminated minus interleaved dark map shows the adduct clearly (+8.6σ; +10.7σ within 1 Å) and no loss of the γ-phosphate (Figure 2B). The laser-off reference is also less isomorphous with the light data (isomorphism R factor R_iso_ 0.096 against 0.041; Table S2). All light-induced changes were therefore computed against the interleaved dark of the same runs, and the laser-off darks were used only to characterize the GDP-bound sub-population.

### The photosensor is activated within 350 µs

In 2026, four light datasets were recorded in the time windows 0–350 µs, 500–850 µs, 1.00–1.35 ms and 1.50–1.85 ms after excitation. A fifth to a quarter of the molecules carry the photoadduct (activated fraction 0.20 in the first window and 0.22–0.24 in the later ones; Table S3). The light-induced difference density is concentrated in the photosensor, and at the chromophore it has the same pattern in all four windows (Figure 3).

**Figure 3.**
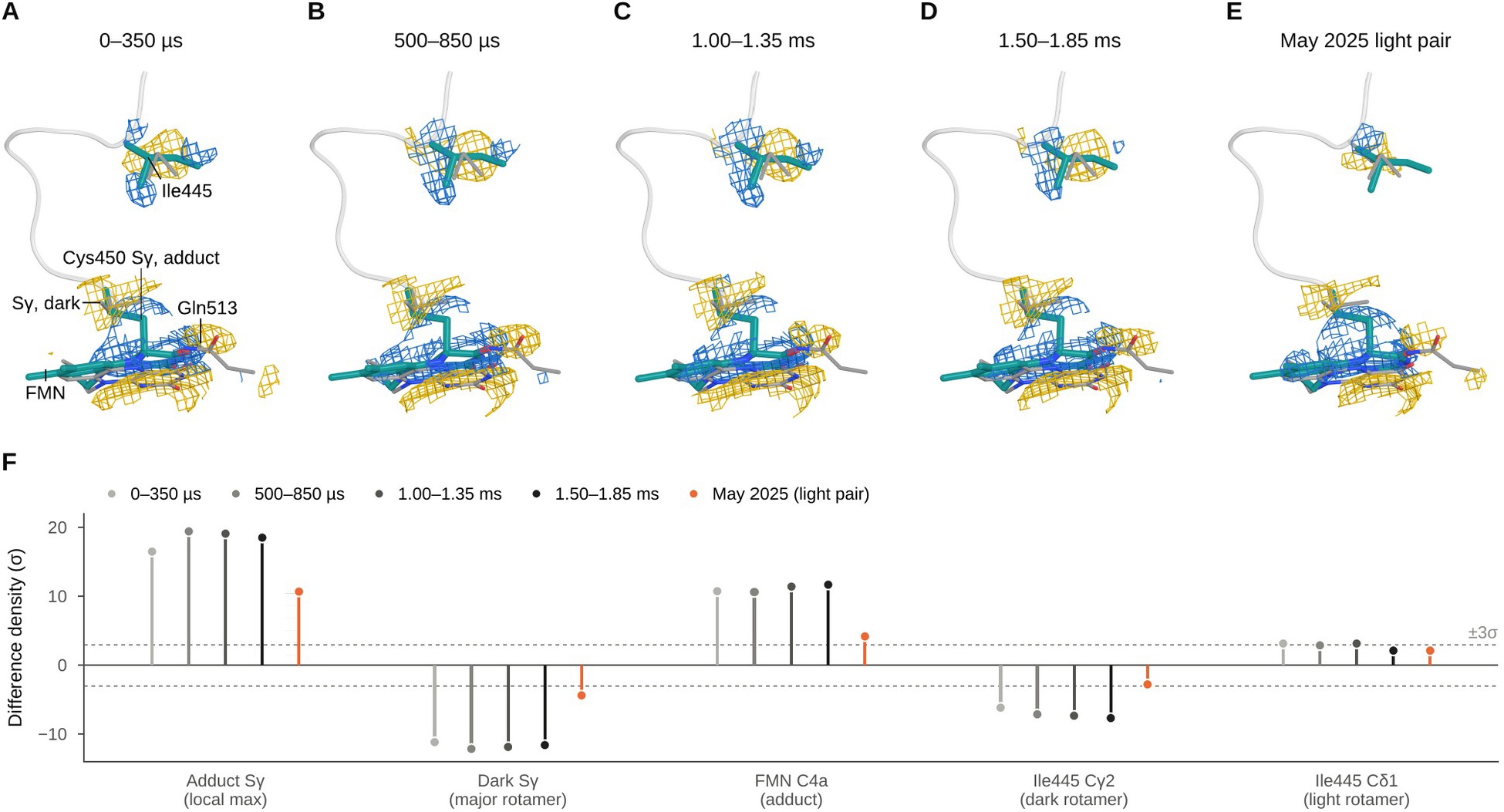
The photosensor is activated within 350 µs. (A–D) Xtrapol8 difference maps (each time window minus the 2026 interleaved dark) at the flavin chromophore for 0–350 µs, 500–850 µs, 1.00–1.35 ms and 1.50–1.85 ms and (E) the 2025 illuminated minus interleaved dark map, contoured at ±3.0σ and carved 1.5 Å around the atoms shown; positive density in blue, negative in gold. Thin gray sticks, dark model (isoalloxazine ring, Cys450 in both rotamers, Ile445 and Gln513); teal sticks, light-state model of each window (adduct-bound Cys450 and flavin, Ile445); gray tube, main chain of residues 444–451. Models are shown as refined, without superposition; for the 2025 map, the 2026 adduct and Ile445 are placed on the 2025 dark model by superposition on LOV2. Asn482, Asn492 and Gln454 are omitted for clarity. (F) Map values at the adduct Sγ (local maximum within 1 Å), the major dark Sγ rotamer, FMN C4a and the dark and light rotamers of Ile445 in each dataset (Table S4); dashed lines, ±3σ.

At the chromophore, the strongest positive feature in every map lies at the position of the covalent Cys450 Sγ atom (+16.5 to +19.4σ), paired with negative density at the Sγ position of the major dark rotamer (−11.1 to −12.1σ), and positive density covers C4a of the adduct-bound flavin (+10.6 to +11.7σ; Table S4). Ile445, which directly follows helix Dα next to Cys450, switches rotamer: the Cγ2 position of its dark rotamer carries −6.1 to −7.6σ, and omit density in every window supports the rotamer with χ1 near −65° that Ile445 adopts in the cryotrapped light-state structures of isolated AsLOV2 (PDB entries 2V0W and 7PGY; Halavaty & Moffat, 2007; Dietler et al., 2022) but not in their dark-state counterparts (2V0U and 7PGX). The features of the first window, in which the pump pulse falls within the first detector frame, are about 15% weaker, in proportion to its lower activated fraction; from 500 µs onward the response does not change.

The residues that hydrogen-bond the flavin (Figure 1E) carry difference density in every window: Asn482 at its side chain (−7.8 to −8.3σ within 1 Å of ND2), Gln454 along its side chain (+5.3 to +6.0σ), Asn492 in strand Hβ on its backbone (down to −5.7 to −6.4σ) and Gln513 at its NE2 atom (down to −4.2 to −5.1σ within 1 Å). In the light-state models, the Gln513 NE2–O4 distance lengthens from 3.14 Å to 3.5–4.2 Å; the orientation of its amide cannot be determined at this resolution and was kept as in the dark state.

The 2025 experiment, excited with 3 ns pulses instead of the 66 µs pulses of 2026, reproduces these changes at an activated fraction of about 0.07–0.09. Against the interleaved dark of the same runs, the merged illuminated data show the adduct (+10.7σ) and negative density at the dark Sγ position (−4.3σ), and the LOV2 features of the 2025 map recur with the same sign in the 2026 maps (Figure S1). The Ile445 rotamer switch stays below 3σ in the difference map at this activation (Table S4), but a polder omit map of the extrapolated data supports the light-state rotamer (Methods).

### The activated photosensor tilts as a rigid body on Rac1

Beyond the chromophore, much of the LOV2 difference density has the pattern of a small rigid-body displacement. Because only LOV2 was refined in the light-state models, we determined this motion from the maps, by fitting them with maps calculated for rigid-body motions of LOV2 and Rac1 (Methods). In every window, a model in which LOV2 rotates relative to Rac1 explains the map substantially better than a single rigid body (R² of 0.25–0.33 against 0.12–0.17), with a tilt of 0.21–0.24° (Table 3), 31 to 38 standard deviations above null maps with randomized signs (Figure S2). Per activated molecule, this is 0.97–1.04°, the same in all four windows within error (Figure 4). The rotation axis is the same in every window (pairwise cosines 0.994–1.000) and passes within 1.3 Å of interface atoms, so the photosensor hinges on Rac1 at the interface. The two 2026 dark datasets give no significant tilt (0.023°).

**Figure 4.**
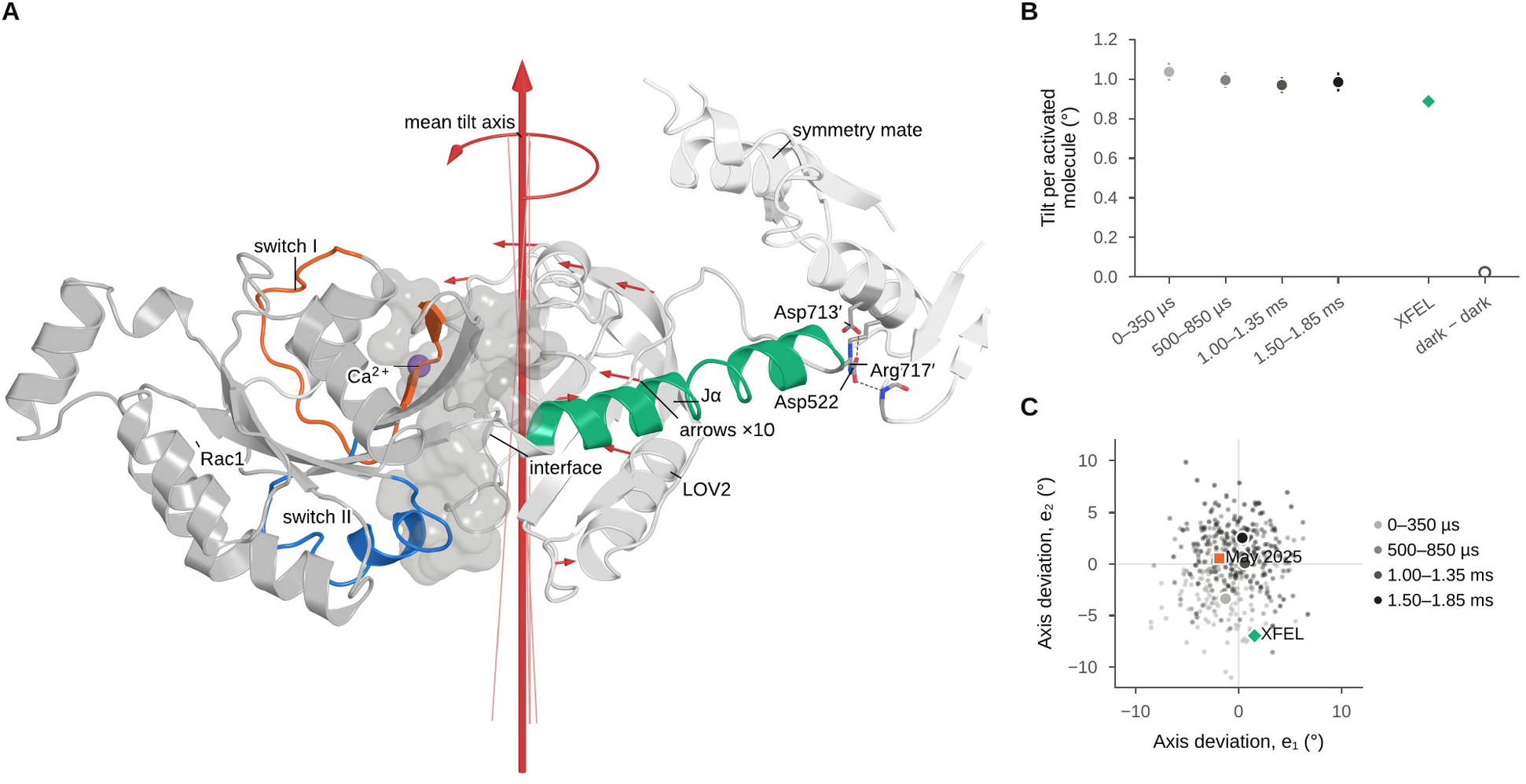
The activated photosensor tilts on Rac1. (A) Tilt of LOV2 relative to Rac1 on the dark-state structure (2026 interleaved dark). Red cylinder, mean axis of the four 2026 windows (thin lines, individual windows, 0.6–3.6° from the mean); curved arrow, sense of the rotation of LOV2 relative to Rac1 (right-hand rule about the axis); straight arrows, Cα displacements of LOV2 for a tilt of 1.0° per activated molecule, exaggerated ten-fold (0.22–0.44 Å before exaggeration). Gray surface, interface atoms (within 4 Å of the other domain); purple sphere, interface Ca²⁺ ion, 15 Å from the axis. The axis passes through the domain interface, 33–35 Å from the lattice contact of Asp522 in Jα with Asp713′ and Arg717′ of a symmetry mate (white cartoon). (B) Tilt per activated molecule in the four 2026 windows (bootstrap standard deviation), with the value for the XFEL light state of the same crystal form (Pachota et al., 2026) and the 2026 dark–dark control. (C) Directions of the fitted tilt axes for the 2026 windows, the 2025 data and the XFEL light state, as angular deviations from the mean 2026 axis; small dots, bootstrap replicates of the 2026 windows.

**Table 3.** Rigid-body tilt of LOV2 relative to Rac1 fitted to the MicroMAX difference maps (2.5–8 Å; Methods). Tilt with bootstrap standard deviation; z, separation from sign-flip null maps in standard deviations; axis cosine with the tilt axis of the XFEL light state of the same crystal form (Pachota et al., 2026); axis distance, distance from the tilt axis to the nearest interface atom (Methods). Activated fractions are from the chromophore-pocket regression (Methods; Table S3). For 2025, phases and fitted model are from the 2025 laser-off dark structure, and values in parentheses use the second estimate of the activated fraction (Methods; Table S3).

| Map | Activated fraction | Tilt (°) | Tilt per activated molecule (°) | z | Axis cosine with XFEL light | Axis distance (Å) | R<sup>2</sup> one body / two bodies |
| --- | --- | --- | --- | --- | --- | --- | --- |

|  |  |  |  |  | state |  |  |
| --- | --- | --- | --- | --- | --- | --- | --- |
| 0–350 μs –<br>interleaved<br>dark | 0.20 | $0.205 \pm$<br>0.009 | 1.04 | 31 | 0.997 | 1.3 | 0.12 /<br>0.25 |
| 500–850 μs –<br>interleaved<br>dark | 0.24 | $0.241 \pm$<br>0.009 | 0.99 | 38 | 0.991 | 0.8 | 0.17 /<br>0.33 |
| 1.00–1.35 ms –<br>interleaved<br>dark | 0.23 | $0.224 \pm$<br>0.010 | 0.97 | 37 | 0.992 | 0.8 | 0.16 /<br>0.33 |
| 1.50–1.85 ms –<br>interleaved<br>dark | 0.22 | $0.214 \pm$<br>0.011 | 0.98 | 35 | 0.986 | 1.0 | 0.16 /<br>0.32 |
| 2025<br>illuminated –<br>interleaved<br>dark | 0.09<br>(0.07) | $0.097 \pm$<br>0.008 | 1.06<br>(1.47) | 20 | 0.990 | 0.3 | 0.03 /<br>0.08 |
| 2026<br>interleaved<br>dark – laser-off<br>dark (control) | – | $0.023 \pm$<br>0.009 | – | 2.6 | 0.63 | 1.8 | 0.03 /<br>0.04 |

The same tilt is present in the 2025 data (0.097 ± 0.008°, 20 standard deviations above null maps; Table 3), about an axis nearly parallel to those of 2026 (cosines 0.998–0.999) that passes 0.3 Å from the nearest interface atom, and of similar size per activated molecule (1.1–1.5°), although the 2025 activated fraction is less well determined. In the light state of the same crystal form accumulated by repeated excitation at an XFEL (XFEL light state; Pachota et al., 2026), LOV2 tilts about the same axis by 0.89° per activated molecule while the Jα helix remains docked (cosines 0.986–0.997 with the axes fitted to the 2026 windows).

The Jα helix itself stays folded and docked in all four windows, up to 1.85 ms: complete disorder of Jα in more than 11–17% of the activated molecules would have been detected (Table S3), although in solution Jα unfolds on a sub-millisecond timescale (Konold et al., 2016, 2024; Kim et al., 2024). In the crystal, the N-terminal end of Jα contacts Rac1 of a symmetry-related molecule (Asp522 with Asp713, Arg717 and the Asp590 amide; Arg521 with Glu714), 33–35 Å from the hinge axis, a contact that may contribute to keeping Jα docked.

### The effector keeps GTP and the dark-state switches

Inside Rac1, the nucleotide-binding site shows no lasting response (**Figure 5**). The γ-phosphorus reads −3.1σ in the first window and +0.3 to +0.8σ thereafter, its three oxygens stay within ±1.4σ and the Mg²⁺ ion within ±0.8σ (Table S4), against −21.8σ at the γ-phosphorus for the GDP-bound sub-population. The first-window value, at the 3σ detection limit (loss in about 3% of the molecules), does not persist into the later windows, as an irreversible loss of the γ-phosphate would. From 500 µs onward, loss of the γ-phosphate in more than 14–18% of the activated molecules is excluded in each window (Table S3). In the light-state models, whose Rac1 coordinates are those of the per-window refinements (Methods), switch I keeps its dark-state geometry: the distance from Thr578 Cβ to the Ala602 carbonyl carbon is 5.96–6.12 Å (6.03 Å in the dark model) and that between the Cα atoms of Ala556 and Tyr575 7.47–7.53 Å (7.52 Å), close to the values of wild-type Rac1 bound to the GTP analog GppNHp (5.82 and 8.24 Å; PDB entry 3TH5; Krauthammer *et al*., 2012) and far from those of GDP-bound Rac1 (see below). Within 1.85 ms of excitation, the nucleotide and the residues that sense the γ-phosphate therefore remain in their dark-state configuration.

**Figure 5.**
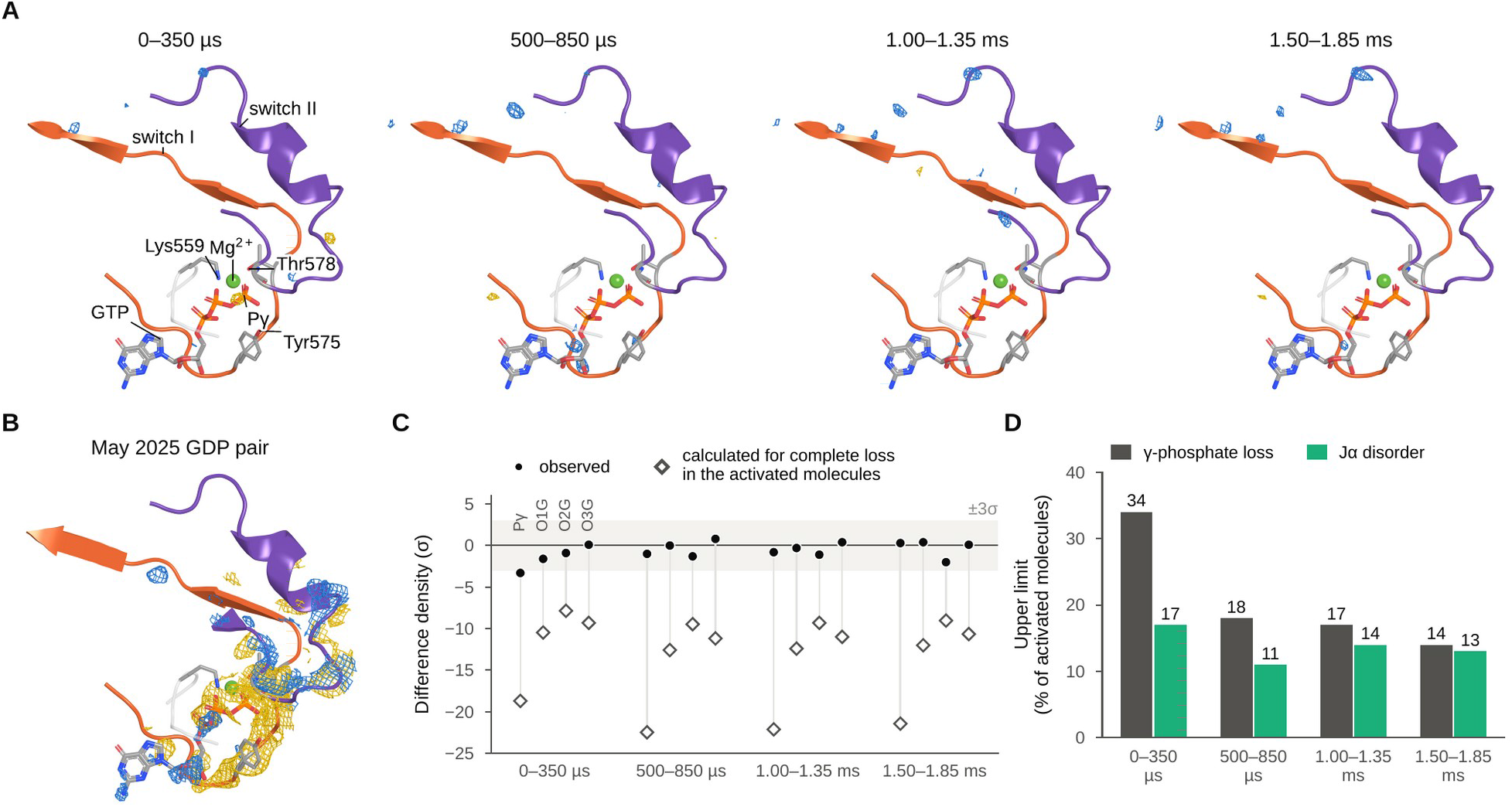
The Rac1 nucleotide site does not respond. (A) Xtrapol8 difference maps (each window minus the 2026 interleaved dark) at the Rac1 nucleotide-binding site and the two switches, contoured at ±3.0σ and carved 1.6 Å around GTP, Mg²⁺ and the main chain of both switches (positive blue, negative gold). Switch I (residues 569–588) is orange, switch II (600–619) purple and the P-loop a gray tube; GTP, Mg²⁺, the side chains of Lys559, Tyr575 and Thr578 and the main chain of Gly603, which contact the γ-phosphate, are shown on the 2026 dark model. (B) The same view for the 2025 interleaved dark minus laser-off dark map, which shows the loss of the γ-phosphate in the GDP-bound sub-population (2025 laser-off dark model). (C) Values at the γ-phosphate atoms in each window, read in the unweighted difference maps used for the detection limits (filled circles; Methods), with the values calculated for complete loss of the γ-phosphate in the activated molecules (open diamonds); shaded band, ±3σ. (D) Upper limits per activated molecule for γ-phosphate loss and Jα disorder (Table S3).

On the Rac1 side, a positive feature recurs in all four windows at the domain interface: it is the strongest positive extremum near switch I in every window, and the strongest in Rac1 in two of them, centered 1.0–1.5 Å from the Ca²⁺ ion (+4.2 to +5.2σ), and it is also present, more weakly, in the 2025 light pair, which has its own dark reference (+2.8σ; Table S4). It therefore reflects a light-induced change at the interface about which LOV2 tilts, although the refined ion moves by only about 0.2 Å, within coordinate error, so we do not assign it to a specific atom. A negative feature at the Gln617–Thr618 peptide (Rac1 Gln74–Thr75), where switch II meets the photosensor, also recurs but stays below 3σ, and features of similar size lie at the lattice contact of Jα (Arg717) and at the modification of Cys648.

### Structure of the GDP-bound sub-population

The GDP-bound sub-population is largest in the 2025 interleaved dark data, which differ from the 2025 laser-off dark at the nucleotide but not at the photosensor. We therefore determined its structure from dark data alone, by extrapolation of the interleaved dark minus laser-off difference (occupancy 0.25; Methods), so that neither light nor a time after excitation is involved.

The difference map is concentrated at a single site (Figure 5B). Its strongest feature is a negative peak of −21.8σ at the γ-phosphorus, with −7.4 to −10.7σ at the γ-phosphate oxygens, while the photosensor shows no difference density (−0.7σ at the adduct Sγ position, +2.1σ within 1 Å). Switch I carries negative density along the segment that binds the γ-phosphate and the Mg²⁺ ion (Tyr575–Thr578, Rac1 Tyr32–Thr35; down to −8.9σ at the Pro577 amide), and switch II carries paired features at Gly603–Asp606 (Rac1 Gly60–Asp63; up to +6.3σ at Gly603 O). A further positive feature lies at the Cys648 modification (+6.7σ).

The refined GDP-state model superposes on the 2025 dark model with an RMSD of 0.47 Å over all 320 Cα atoms, and the two domains move as one unit: whichever domain is used for the fit, the other superposes at 0.21–0.25 Å when switch I (residues 570–579) is excluded, and Jα remains folded and docked.

The largest change lies in switch I. After superposition on the Rac1 core, Gly573 (Rac1 Gly30) is displaced by 3.9 Å, Glu574–Thr578 (Rac1 Glu31–Thr35) by 2.7–3.4 Å and Val579 and Phe580 by only 0.5 and 0.2 Å (Figure 6A, C), whereas switch II moves by at most 0.7 Å. Switch I remains mobile (main-chain B factors 99–137 Å²) but follows a defined path: in a polder omit map with residues 569–583 omitted, its main-chain density is 3.5–4.3σ for Glu574–Thr578 and 3.8–6.7σ for Val579–Tyr583, against 4.1–6.8σ for a Rac1 core segment omitted in the same way, and weaker only at Pro572 and Gly573 (2.9 and 1.9σ; Figure S3). The 2025 illuminated data referenced to the laser-off dark, which contain the same sub-population in independent images, and a re-refinement without switch I give the same result. Switch I is thus released from the nucleotide into a mobile, partially open conformation whose main-chain path is defined from Glu574 to Tyr583, although that of Pro572–Gly573 and the side chains are not.

**Figure 6.**
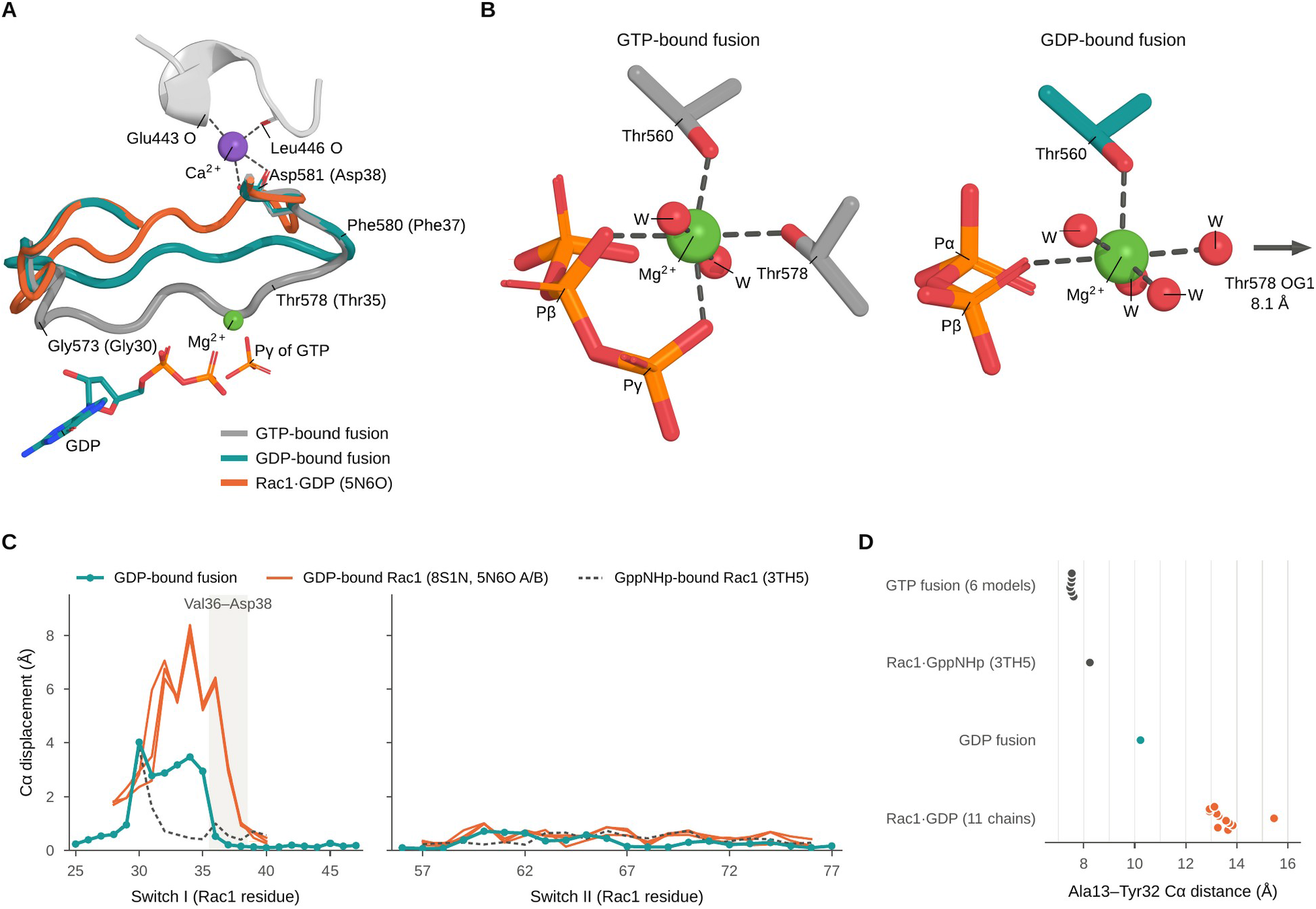
Structure of the GDP-bound sub-population. (A) Switch I (residues 567–584, Rac1 24–41) of the GTP-bound fusion (2025 laser-off dark model, gray), the GDP-bound fusion (GDP-state model, teal) and free GDP-bound Rac1 (PDB 5N6O chain A, orange) after superposition on the Rac1 core of the GTP-bound fusion, drawn as tubes through the Cα atoms. GDP and Mg²⁺ of the GDP-state model, the γ-phosphate of GTP in the GTP-bound fusion, and the interface Ca²⁺ ion (purple) with its ligands Asp581 (Rac1 Asp38) and the Glu443 and Leu446 carbonyl oxygens of LOV2 (gray cartoon) are shown. (B) Mg²⁺ coordination in the GTP-bound fusion (2025 laser-off dark model) and in the GDP-bound fusion (GDP-state model), viewed close to the axis through two trans water molecules (W). In the GDP-bound fusion, Thr578 (Rac1 Thr35) OG1 lies 8.1 Å from the ion (arrow), and four water molecules complete the coordination with Thr560 OG1 and a β-phosphate oxygen (Mg²⁺–ligand distances 2.05–2.20 Å; Table S5). (C) Cα displacement of switch I and switch II from the GTP-bound fusion after superposition on the Rac1 core, for the GDP-bound fusion (teal), free GDP-bound Rac1 (8S1N, 5N6O; orange) and GppNHp-bound Rac1 (3TH5; dashed); the band marks Val36–Asp38, next to the interface Ca²⁺ ion. (D) Ala13–Tyr32 Cα distance in the GTP-bound models of the fusion, GppNHp-bound Rac1, the GDP-bound fusion and the eleven Rac1 chains of six deposited GDP-bound structures (orange).

Every contact to the γ-phosphate present in the dark state is absent in the GDP-state model, from Thr578 OG1 and its amide, Tyr575 OH, the Gly603 amide, Lys559 NZ, the Mg²⁺ ion and three water molecules (Table S1), and Lys559 NZ keeps its contact to the β-phosphate (2.62 Å). Thr578 (Rac1 Thr35) leaves the Mg²⁺ coordination sphere, its OG1 now 8.1 Å from the ion, which is octahedrally coordinated by Thr560 OG1 (2.12 Å), a β-phosphate oxygen (2.20 Å) and four water molecules (2.05–2.17 Å; Figure 6B; Table S5). The P-loop and guanine contacts are kept, so the nucleotide stays in its pocket and the change is confined to the phosphate end that the switches read. How the γ-phosphate is lost from a hydrolysis-deficient Q61L construct is considered in the Discussion.

We compared the GDP-bound fusion with six deposited structures of GDP-bound Rac1, two of free Rac1 (PDB entries 8S1N and 5N6O; Dubois *et al*., 2026; Ferrandez *et al*., 2017), three of complexes with RhoGDI, arfaptin and the *Yersinia* effector YpkA (1HH4, 1I4D and 2H7V; Grizot *et al*., 2001; Tarricone *et al*., 2001; Prehna *et al*., 2006) and one of a Zn²⁺-bound trimer (2P2L; Prehna & Stebbins, 2007), and with GppNHp-bound Rac1 (3TH5), after superposition on the Rac1 core. In free GDP-bound Rac1, switch I is displaced from its position in the GTP-bound fusion by 5.2–8.4 Å over Tyr32–Val36 (Rac1 numbering), whereas in the GDP-bound fusion the displacement is at most 3.9 Å and falls to 0.5 Å at Val36 and 0.2 Å at Phe37 (Figure 6C). The part of switch I that stays in place lies at the interface with the photosensor, where Asp581 (Rac1 Asp38) keeps coordinating the Ca²⁺ ion with both carboxylate oxygens (2.58 and 2.64 Å). Switch II does not distinguish the states: its Cα atoms lie within 1.5 Å of their positions in the GTP-bound fusion in GppNHp-bound Rac1, in the GDP-bound fusion and in all deposited GDP-bound structures except the complexes 2H7V and 2P2L.

The Cα distance between Ala556 in the P-loop and Tyr575 in switch I (Rac1 Ala13 and Tyr32) summarizes this difference (Figure 6D): it is 7.47–7.65 Å in the GTP-bound models of the fusion, 8.24 Å in GppNHp-bound Rac1, 10.20 Å in the GDP-bound fusion and 12.93–15.46 Å in all eleven Rac1 chains of the deposited GDP-bound structures. Our data do not support the switch I conformation of free GDP-bound Rac1: placed by the core superposition, its Cα atoms of Gly30–Phe37 lie in mean polder density of 1.6–1.7σ, against 3.7σ for the GDP-state model, and carry no more density than random placements of the same atoms once the modeled switch I is accounted for (Figure S3B; Methods). Because Gly573–Thr578 is mobile, we do not interpret its precise conformation, but switch I is less open in the GDP-bound fusion than in GDP-bound Rac1 alone, and the difference lies where switch I meets the photosensor.

### A partially open switch I can persist in simulations of the GDP-bound fusion

To test whether this conformation requires the crystal lattice, we simulated the GDP-bound fusion in explicit water (Methods). Starting from the 0–350 µs structure with GDP in place of GTP, the Mg²⁺ ion adopted the coordination of the GDP-state model (Table S5). Adaptive biasing of selected distances and angles brought the switches to the geometry of an earlier model of the GDP-bound sub-population (Methods), which agrees with the GDP-state model to 0.44 Å (switch I) and 0.15 Å (switch II) Cα; the restraints were then released gradually and the fusion was followed for 40 ns without bias. In two independently seeded trajectories, both switches stayed close to this target, with transient excursions (Figure 7): over the final 5 ns, the median Cα RMSDs from the target were 1.01 and 1.03 Å for switch I and 1.30 and 1.11 Å for switch II, and 97.8–99.0% of frames lay below the 95th percentile of the RMSDs in a simulation started from the target model itself. The GDP-like Mg²⁺ coordination shell persisted in every saved frame, and the interface Ca²⁺ ion stayed coordinated by Asp581 (Rac1 Asp38) throughout both trajectories, whereas the Glu443 carbonyl left its coordination sphere.

**Figure 7.**
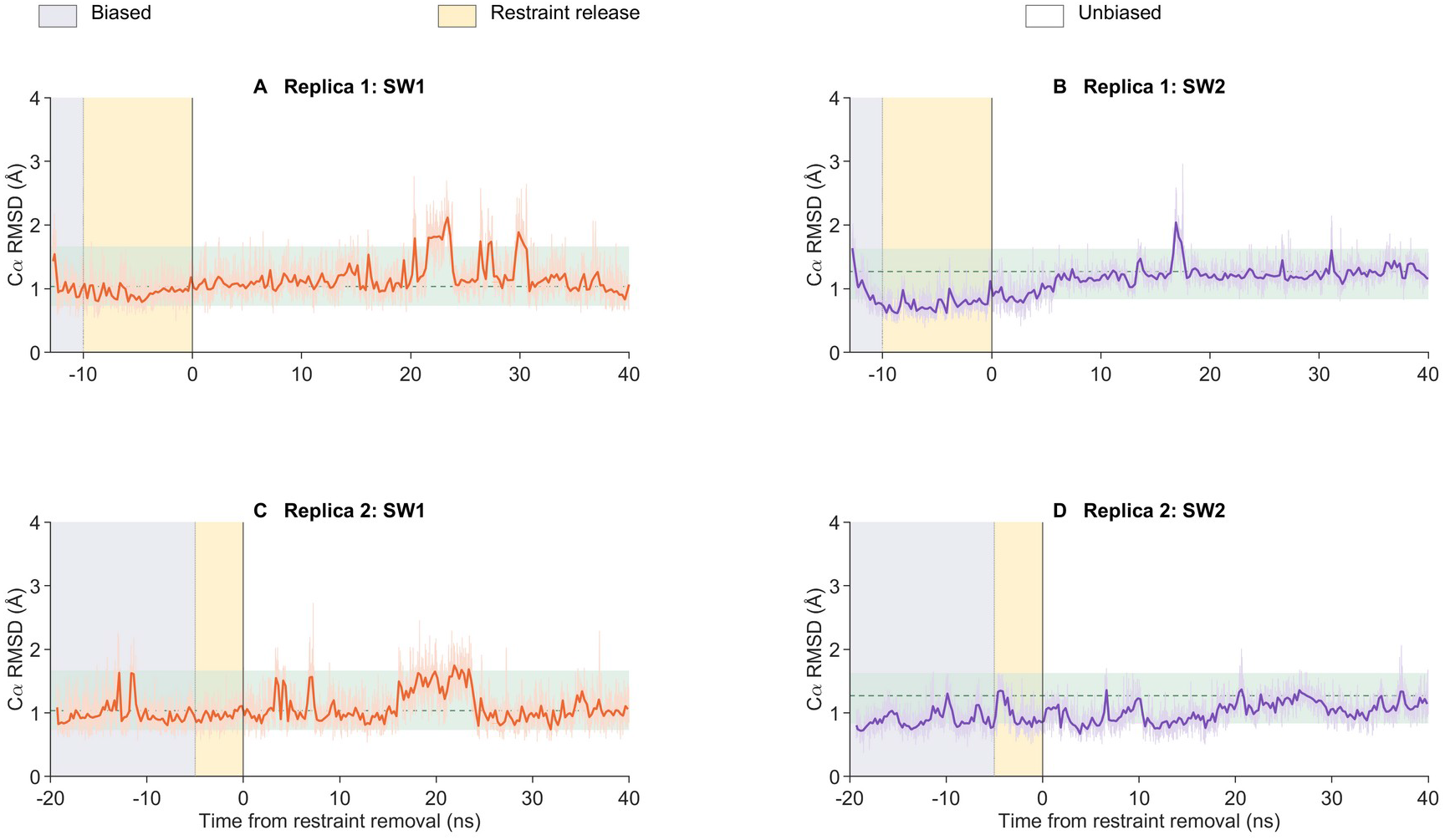
Switch geometry of the GDP-bound fusion in molecular dynamics simulations. Cα RMSDs of switch I (SW1, Rac1 residues 28–40; orange) and switch II (SW2, Rac1 residues 60–76; purple) from an earlier model of the GDP-bound sub-population used as the target (Methods), after superposition on Rac1 residues 4–180 without both switches, for replica 1 (A, B) and replica 2 (C, D). Gray background, biased dynamics; amber, gradual release of the restraints; white, unbiased dynamics. Time zero marks complete removal of the restraints. Thin traces show saved frames and thick traces medians in non-overlapping 0.25 ns bins. Green bands and dashed lines show the 5th–95th percentile range and the median of a simulation started from that model.

Not all biased trajectories retained this geometry after the restraints were released, and the simulations do not describe how or how fast switch I rearranges after loss of the γ-phosphate. They show that a partially open conformation like that of the crystal structure is accessible to the GDP-bound fusion and can persist for tens of nanoseconds without lattice contacts.

## Discussion

The 2026 time series shows that, in the crystal, the photosensor of PA-Rac1 is activated before the effector responds. Within 350 µs of excitation, about a fifth of the molecules carry the photoadduct, Ile445 has switched rotamer and LOV2 has tilted on Rac1; these changes then grow only with the activated fraction, while GTP, the Mg²⁺ ion and both switches of Rac1 keep their dark-state configuration up to 1.85 ms. The response of the photosensor to light is thus established within 350 µs, consistent with microsecond adduct formation and β-sheet changes in solution (Swartz *et al*., 2001; Gil *et al*., 2017), and within 1.85 ms it reaches the effector only as a rigid-body motion at the shared interface.

In solution, Jα in AsLOV2 unfolds on a sub-millisecond timescale (Konold *et al*., 2016, 2024; Kim et al., 2024), and in PA-Rac1 illumination shifts the population of pre-existing caged and uncaged conformations rather than releasing Rac1 (Winkler *et al*., 2015). In the crystal, Jα stays docked, as in the XFEL light state of the same crystal form, and its N-terminal end forms a lattice contact. The state we observe, with the adduct formed, Ile445 switched and Jα docked, may correspond to the intermediate that solution scattering resolves after adduct formation and before Jα unfolding (Konold *et al*., 2024). In this state, the whole photosensor tilts on Rac1 about an axis through the domain interface, by a similar amount per activated molecule at a synchrotron and at an XFEL. We propose that this hinge motion is the crystalline counterpart of the first step of uncaging, a loosening of the LOV2–Rac1 interface that in solution would precede or accompany the release of Jα. The feature that recurs in all four windows on the Rac1 side lies at this interface, next to the Ca²⁺ ion that bridges the two domains, so the first change we detect on the effector side is at the interface itself, not in the switches or at the nucleotide.

The GDP-bound sub-population shows how Rac1 within the fusion accommodates the loss of the γ-phosphate. Thr578 leaves the Mg²⁺ coordination sphere and the part of switch I that binds the γ-phosphate and the Mg²⁺ ion is released, whereas switch II and the guanine base stay in place, as expected from the canonical switch mechanism of small GTPases (Vetter & Wittinghofer, 2001). Switch I, however, opens only partially: by at most 3.9 Å, against 5–8 Å from Tyr32 to Val36 (Rac1 numbering) in GDP-bound Rac1 alone, with Val36 and Phe37, next to the Ca²⁺ ligand Asp38, in place. Switch II stays close to its GTP-bound position also in GDP-bound Rac1 alone, so its retention in the fusion does not by itself indicate a restraint by the photosensor.

We propose that the docked photosensor restrains switch I. The C-terminal end of switch I is held at the domain interface through the Ca²⁺ ion coordinated by Asp581 (Rac1 Asp38) and by the main-chain carbonyls of Glu443 and Leu446 of LOV2, so that only the segment that binds the γ-phosphate and the Mg²⁺ ion can follow the nucleotide; without lattice contacts, this partially open conformation, once reached, persisted for 40 ns in simulations. This is the interface about which the activated photosensor tilts and at which a light-induced feature recurs on the Rac1 side, so the LOV2–Rac1 interface is where photosensor and GTPase meet in both situations we observe, when the photosensor is activated and when the GTPase loses its γ-phosphate. In our crystals this interface is bridged by the Ca²⁺ ion, which holds the C-terminal end of switch I in the GDP-bound state. The ion is found in our crystals and in 2WKP, both grown with calcium acetate, but not in 2WKQ and 2WKR, grown without it (Wu et al., 2009). Ca²⁺ also favors the caged conformation of PA-Rac1 in solution (Winkler et al., 2015), but whether the site is occupied at cellular Ca²⁺ concentrations is not known, so we do not assign the ion a role in signaling. Structures and simulations of the fusion without calcium or without the LOV2 domain, and time-resolved structures at longer delays, can test whether the photosensor restrains switch I and whether this restraint is relieved when Jα unfolds. If it is, the photosensor of PA-Rac1 acts not only by occluding the effector-binding surface but also on the conformation of the GTPase itself, as in Cdc42Lov, an engineered fusion of LOV2 and Cdc42 in which NMR shows light-state perturbations that reach the effector-binding site and coupling between the two domains in both directions (Jain *et al*., 2023).

How the γ-phosphate is lost remains open. It is lost in the crystals without illumination, to an extent that differs between samples, and light does not drive the loss on the timescales probed (Table 2; Pachota et al., 2026). Because Q61L Rac1 does not hydrolyze GTP measurably in solution (Davis *et al*., 2013), slow attack of the water that lies 3.5 Å from the γ-phosphorus, close to in line with the scissile bond and with no side chain positioned to act as a base (Figure 1D), is a plausible route.

These observations are relevant to the use of PA-Rac1 as an optogenetic tool. A GDP-bound fraction of the fusion would remain inactive even when the photosensor releases the effector-binding surface, and because the E91H and N92H substitutions diminish interactions with regulatory proteins (Wu *et al*., 2009), such a fraction might be reloaded with GTP only slowly. Whether this occurs in cells and limits the dynamic range of the tool is not known; it can be tested by measuring GTP hydrolysis by the fusion in the dark and under continuous blue light, with the light-insensitive C450A variant as a control.

In summary, time-resolved crystallography of an engineered photoswitch shows its photosensor activated within 350 µs and tilting on an unchanged GTP-bound effector about their shared interface, and a GDP-bound state shows switch I following the loss of the γ-phosphate only partially, held at a Ca²⁺ ion at the same interface. Referencing the pump–probe data to interleaved dark frames was essential for both conclusions, because the dark samples themselves differ in their nucleotide content. For LOV2-based tools, these results point to the interface between photosensor and effector as a feature to consider alongside the Jα linker when caging of the effector is optimized.

### Limitations of the study

Jα stayed docked within 1.85 ms in the crystal, whereas it unfolds on a sub-millisecond timescale in solution; its N-terminal end forms a lattice contact that may contribute to this, so the structures may represent an early stage of the photosensor response. As is usual in time-resolved crystallography, a fraction of the molecules was activated, about a fifth here, and the light states were therefore characterized from extrapolated amplitudes and difference maps; within these data, loss of the γ-phosphate in fewer than 14–18% of the activated molecules would remain below the detection limit. In the light-state models LOV2 was refined, and the relative orientation of the two domains was determined directly from the difference maps. The GDP-bound sub-population forms without illumination by a route we have not established, and its switch I is defined by the density from Glu574 onward. The simulations reached the partially open conformation under bias, and the activity of the fusion was not measured, so the functional consequences we propose remain to be tested.

## Resource availability

### Lead contact

Requests for further information and resources should be directed to and will be fulfilled by the lead contact, Przemyslaw Nogly.

### Materials availability

The expression plasmid of PA-Rac1 used in this study is available from the lead contact upon request.

### Data and code availability

- Coordinates and structure factors have been deposited in the Protein Data Bank under accession codes 35EU (2025 laser-off dark), 35EV (2025 light state), 35FB (2025 GDP state), 35FI (2026 interleaved dark) and 35FN, 35FK, 35FL and 35FT (2026 light states at 0–350 µs, 500–850 µs, 1.00–1.35 ms and 1.50–1.85 ms) and are publicly available as of the date of publication. The structure-factor files of the extrapolated datasets also contain the merged intensities of the triggered and reference datasets (Methods). Raw diffraction images are available from the lead contact upon request.
- The molecular dynamics input files and trajectories will be deposited at Zenodo before publication; until then, they are available from the lead contact upon request.
- Any additional information required to reanalyze the data reported in this paper is available from the lead contact upon request.

## Acknowledgments

We acknowledge MAX IV Laboratory for time on beamline MicroMAX under proposals 20241674 (2025) and 20251858 (2026). Research conducted at MAX IV, a Swedish national user facility, is supported by the Swedish Research Council under contract 2018-07152, the Swedish Governmental Agency for Innovation Systems under contract 2018-04969 and Formas under contract 2019-02496. MicroMAX is funded by the Novo Nordisk Foundation under the grant number NNF17CC0030666. We thank Bruce Doak for providing the high-viscosity extrusion injector designed for MicroMAX. We acknowledge the P11 beamline of PETRA III operated by DESY Photon Science (No. R-20240820 EC) for cryo-crystallography beamtime carried out in preparation for these experiments. We acknowledge the European Synchrotron Radiation Facility (ESRF) for provision of synchrotron radiation facilities under proposals MX-2593 and MX-2692, and we thank the beamline staff for assistance and support. We acknowledge the Malopolska Centre of Biotechnology Structural Biology Core Facility for access and support. We thank the Department of Microbiology, Faculty of Biochemistry, Biophysics and Biotechnology, Jagiellonian University, for access to its bacterial culture and cell disruption infrastructure.

The project was financed under Dioscuri, a program initiated by the Max Planck Society, jointly managed with the National Science Centre in Poland, and mutually funded by the Polish Ministry of Science and Higher Education and the German Federal Ministry of Research, Technology and Space. This research was funded by the Polish National Science Centre (grant agreement No. UMO-2021/03/H/NZ1/00002 and SONATA BIS grant No. UMO-2024/54/E/NZ1/00429, both awarded to P.N.). The purchase of equipment used in the project has been supported by a grant of the Faculty of Biochemistry, Biophysics and Biotechnology under the Strategic Programme Excellence Initiative at Jagiellonian University. The work presented in this paper benefits from the support received from the Polish Ministry of Science and Higher Education, dec. no. 2021/WK/11. E.R. acknowledges support from EPSRC grant EP/R013012/2 (Novel Enhanced Sampling Methods in Multiscale Modeling), HECBioSim (EP/X035603/1) and CCPBioSim (EP/T026308/1).

## Author contributions

Conceptualization and coordination: P.N. Methodology: M.P., M.K.S., C.M.C., J.N., P.N. Investigation (protein production and crystallization): M.P., A.K., A.S., A.A., R.K. Investigation (sample delivery): A.K., A.S., M.P. Investigation (TR-SSX data collection at MicroMAX): S.M.K., L.L.C., M.C., C.M.C., D.L., M.B., M.M., J.N., M.P., M.K.S., M.W., B.O., A.K., A.S. Formal analysis (data processing, difference maps and refinement): C.M.C., M.K.S., B.O., P.N. Formal analysis (molecular dynamics): E.R. Resources: T.U., J.N. Visualization: P.N., M.P., M.K.S., E.R. Writing, original draft: P.N., E.R., M.K.S. Writing, review and editing: C.M.C., M.B., M.P., S.M.K., M.C., T.U., J.N., M.K.S., E.R., P.N. Supervision: P.N., J.N., E.R. Funding acquisition: P.N., E.R.

## Declaration of generative AI and AI-assisted technologies in the writing process

During the preparation of this work, the authors used Claude (Anthropic) and Codex (OpenAI) to write and run analysis and figure scripts, to assist with software development and to draft and edit the text. All methods, analyses, results and conclusions were designed, verified and approved by the authors. After using these tools, the authors reviewed and edited the content as needed and take full responsibility for the content of the publication.

## Supplemental information

Data S1. Atom definitions of the restraints in the molecular dynamics simulations (CSV), related to Figure S5 and Methods.

## Methods

### Protein expression and purification

The expression and purification protocol was based on Wu *et al*. (2009) with modifications. A codon-optimized construct encompassing amino acids 404–546 of *Avena sativa* phototropin 1 and 4–180 of human Rac1 with the Q61L, E91H and N92H substitutions of PA-Rac1 (Wu et al., 2009) and an N-terminal tag (MGGSHHHHHHHHGS), cloned into the pQE-60 vector, was obtained from GenScript. The plasmid was transformed into *Escherichia coli* XL10-Gold and cultures were grown in LB medium until OD_600_ reached 0.6–0.8. Expression was induced with 1 mM isopropyl β-D-thiogalactopyranoside and continued overnight at 27 °C. Cells were harvested by centrifugation (20 min, 6,000 × g, 4 °C) and pellets stored at −80 °C.

Thawed pellets were resuspended in lysis buffer containing 20 mM Tris-HCl pH 8.5, 50 mM NaCl, 5 mM MgCl₂, 2 mM β-mercaptoethanol and 40 µM GTP, and disrupted by a single passage through a continuous-flow cell homogenizer at 23,000 psi (159 MPa). The lysate was clarified by centrifugation (1 h, 30,000 × g, 4 °C) and applied to a HisTrap HP column. The His-tagged protein was eluted over a linear imidazole gradient and dialyzed overnight at 4 °C against 10 mM Tris-HCl pH 8.5, 20 mM NaCl, 5 mM MgCl₂, 2 mM β-mercaptoethanol and 40 µM GTP. The sample was applied to a Capto HiRes Q 5/50 column and eluted over a linear NaCl gradient. Protein-containing fractions were pooled and resolved on a HiLoad Superdex 75 pg column in 10 mM Tris-HCl pH 8.5, 20 mM NaCl, 5 mM MgCl₂, 2 mM DTT and 40 µM GTP. GTP was included in all buffers throughout purification, which yielded a more homogeneous preparation better suited to downstream applications. The protein was concentrated to 18 mg ml⁻¹, aliquoted and flash-frozen in liquid nitrogen.

### Crystallization

The crystallization conditions of Wu *et al*. (2009) (12% (w/v) PEG 8000, 100 mM sodium cacodylate pH 5.5 and 100 mM calcium acetate; PDB entry 2WKP) were first reproduced, after which a batch protocol was developed to produce the crystal quantities required for serial crystallography. Protein at 18 mg ml⁻¹ was added dropwise to a precipitant solution containing 18% (w/v) polyethylene glycol 8000, 100 mM sodium cacodylate pH 5.5 and 150 mM calcium acetate in a 2:1 ratio (protein:precipitant, v/v) and mixed gently. Crystallization vessels were incubated in the dark at 20 °C. Long rod-shaped crystals 20–300 µm in thickness appeared within 24 h and sedimented to the bottom of the vessel.

### Sample preparation for serial synchrotron crystallography

Crystals were sedimented by centrifugation and resuspended in mother liquor at the desired density. The crystal slurry was mixed 1:1 with an 18% (w/v) hydroxyethyl cellulose matrix (**final concentration** 9**% (w/v)**), which additionally fragmented the large needles into microcrystalline pieces and removed the need to grow uniform microcrystals. Resuspended crystals were loaded from the back of a Hamilton syringe and mixed with the hydrated viscous matrix using a three-way syringe coupler (James *et al*., 2019; Gotthard *et al*., 2024b).

### TR-SSX data collection at MicroMAX

We performed two time-resolved serial synchrotron crystallography (TR-SSX) experiments at the MicroMAX beamline of MAX IV (Gonzalez *et al*., 2025), in May 2025 and April 2026. In both, pump–probe data were recorded together with interleaved dark frames from the same sample stream, and a separate dark dataset was recorded with the laser off; both experiments were processed, extrapolated and refined with the same protocol (described below). The sample was delivered with a high-viscosity extruder (Nogly *et al*., 2015; Weinert *et al*., 2019; Shilova et al., 2020) equipped with a 75 µm nozzle and operated at a flow rate of 5.32 µl min⁻¹, producing a stable continuous stream traveling at 20.06 mm s⁻¹ (approximately 20 µm ms⁻¹). In 2025, data were collected at a photon energy of 13.016 keV, with the X-ray beam focused to approximately 15 × 5 µm (horizontal × vertical) at the sample and a photon flux of 7.7 × 10¹s photons s⁻¹; in 2026, at 13.699 keV, with a beam of 20 × 5 µm (horizontal × vertical, FWHM) and 8.6 × 10¹s photons s⁻¹. Diffraction data were collected with the PSI-developed adaptive-gain, charge-integrating JUNGFRAU 9M detector (Mozzanica et al., 2018; Leonarski et al., 2018; Duan et al., 2026).

### 2025 experiment

The laser-off dark dataset was recorded first. Pump–probe data were then recorded with a 450 nm laser delivering pulses of 3 ns duration and about 2 µJ energy, focused to 80 × 80 µm (1/e²) at the sample, at a repetition rate of 50 Hz. Diffraction images were recorded in consecutive frames with a frame time of 500 µs and an integration time of 170 µs, matching the 170 µs X-ray pulse generated by an X-ray chopper (Celeroton AG, Switzerland); the frames recorded before each pump pulse (classes 0–3) were merged into the 2025 interleaved dark dataset and the frames after it (classes 4–7) into the 2025 illuminated dataset. Because the activated fraction was lower than in 2026 (Results), the illuminated classes were merged and used to test the light response of the photosensor without time resolution.

### 2026 experiment: time-resolved series

The intermediate states were recorded with the pump–probe scheme established at MicroMAX (Olasz *et al*., 2026b). The sample was first probed in the dark state, then excited by a 450 nm laser pulse and probed again by the X-rays. Diffraction images were recorded in consecutive frames with a frame time of 500 µs and an integration time of 350 µs, matching the 350 µs X-ray pulse generated by an X-ray chopper (Celeroton AG, Switzerland), which set the shortest accessible time window. The laser pulse was synchronized with the start of a detector frame, so that the four frames following each pulse probe the time windows 0–350 µs, 500–850 µs, 1.00–1.35 ms and 1.50–1.85 ms after excitation (classes 4–7). Frames recorded before each laser pulse were merged to give the interleaved dark reference. A separate dark dataset was recorded with the laser off. Throughout the text, the light datasets are referred to by their time window, and each was analyzed separately. Excitation was provided by 450 nm laser pulses of 66 µs duration and about 3 µJ energy at the sample. The laser beam measured 103 µm (horizontal) × 86 µm (vertical) at the sample position, and the laser operated at a repetition rate of 50 Hz.

At the stream velocity of approximately 20 µm ms⁻¹, the illuminated sample volume moves by about 37 µm by the end of the last time window (1.85 ms) and therefore remains within the laser spot, whereas between successive laser pulses at 50 Hz the stream advances by about 400 µm, so that each pulse excites fresh sample and no crystal is illuminated twice. This is important for PA-Rac1, whose photoadduct decays thermally within about a minute (τ = 68 s for isolated AsLOV2 at 22 °C; Nash *et al*., 2008), because re-excitation of partially recovered crystals would contaminate the dark reference. Frames recorded after the fourth light-activated window, during which the illuminated volume may still pass the X-ray interaction point, were not used in the analysis.

### Data processing

Data were processed using the MicroMAX in-house developed, *CrystFEL*-based (White *et al*., 2012, 2016) data reduction workflow, which includes automated optimization of the beam position and peakfinder8 parameters as well as time-delay classification of frames. Peak search used *peakfinder8* (Barty *et al*., 2014) with --threshold=10, --min-snr=4 and --min-pix-count=2, and indexing used *mosflm* (Powell, 1999) for the 2025 datasets and xgandalf (Gevorkov *et al*., 2019) for the 2026 datasets. The indexing ambiguity inherent to space group P3₂21 was resolved with *ambigator* (White *et al*., 2016), using -y 321_H and --operator=-h,-k,l. Intensities were scaled and merged with *partialator* using the unity partiality model, and data statistics were calculated with *check_hkl* and *compare_hkl*. The resulting *hkl* files were converted to *mtz* format with *f2mtz* from the CCP4 suite (Winn *et al*., 2011). The 2025 and 2026 datasets are indexed in alternative settings related by −h,−k,l; the 2025 data were reindexed before any comparison between the two experiments.

Crystals belonged to space group P3₂21 with unit-cell parameters for 2025/2026 datasets *a* = *b* = 115.44/115.48 Å, *c* = 69.94/70.00 Å, α = β = 90°, γ = 120°. Dataset statistics are reported in Table 1.

### Difference maps and dark references

Difference maps and extrapolated structure-factor amplitudes were calculated with *Xtrapol8* v1.2.9 (De Zitter *et al*., 2022), with identical settings for both experiments. For 2026, each time window was the triggered dataset and the 2026 interleaved dark the reference (2.2 Å); the windows were not merged. For 2025, three pairs of merged datasets were used: illuminated (classes 4–7) minus interleaved dark (classes 0–3; 2.4 Å; the light pair) for the light response, interleaved dark minus laser-off dark (2.6 Å; the GDP pair) for the GDP-bound sub-population (Results), and illuminated minus laser-off dark (2.6 Å) as a reference. Intensities were converted to amplitudes by French–Wilson treatment (French & Wilson, 1978) in cctbx (Grosse-Kunstleve *et al*., 2002), the triggered amplitudes were scaled anisotropically to the reference amplitudes, and amplitude differences were weighted (Ursby & Bourgeois, 1997). Phases were calculated from the dark model of the same experiment (see Model building and refinement). Map values are given in σ units of the whole unit cell, read with *GEMMI* (Wojdyr, 2022) on a grid of sample rate 4 with tricubic interpolation, either at a model atom or as the local extremum within 1 Å of it, as stated.

To compare the dark references, the dark datasets of both experiments, the four interleaved dark classes and the illuminated data of 2025 were refined with an identical protocol in *Servalcat* (Yamashita *et al*., 2021, 2023): French–Wilson treatment of the merged intensities (French & Wilson, 1978), five cycles of restrained refinement with isotropic B factors and without hydrogens, starting from the dark model of the same year and with R-free flags copied from the same-year refinement (2025 datasets at a common resolution of 2.45 Å). The 2026 laser-off data, which had been merged in the 2025 indexing setting, were first reindexed by −h,−k,l. The occupancy of the γ-phosphate (PG, O1G, O2G and O3G) was then refined as a single group, once with coordinates fixed and its B factor tied to that of the β-phosphorus (B(Pβ) + 3 Å²), and once with B factor and coordinates refined freely. Dark-minus-dark difference maps were calculated between 8 and 2.5 Å after scaling in resolution shells, with phases from the refined model of the reference dataset.

### Extrapolation, occupancy and activated fraction

Extrapolated amplitudes were obtained as qFextr = F_ref_ + αq(F_trig_ − F_ref_), where q is the normalized weight and α the inverse of the triggered-state occupancy; negative values were corrected by French–Wilson truncation. The occupancy was set by difference-map maximization in *Xtrapol8*: 0.15 in every 2026 window, 0.08 for the 2025 light pair and 0.25 for both 2025 pairs referenced to the laser-off dark, with the Pearson criterion agreeing within one scan step (Table S2, Figure S4). The residues that Xtrapol8 selected for the occupancy determination were Cys450, FMN and Leu480 (with Ile481 at 0–350 µs) in every 2026 window, with no Rac1 residue. *Xtrapol8* was run with Phenix 1.19.2 (Liebschner *et al*., 2019), with CCP4 SCALEIT and TRUNCATE replaced by their cctbx equivalents. The R-free flags of the refinement each dataset started from were kept (see Model building and refinement).

The *Xtrapol8* occupancy is the value that optimizes the extrapolated map. It underestimates the fraction of activated molecules, because difference features in maps phased with the dark model vanish before the full light state is reached. The activated fraction was therefore estimated separately, by regressing the observed light-minus-dark density on the density calculated for complete adduct formation over grid points of the chromophore pocket, using the amplitude scale of each map and the phases of its dark model (unweighted difference maps, 2.2–8 Å for 2026 and 2.4–8 Å for 2025). For the simulation, Cys450 and FMN of the 0–350 µs model were placed in the dark model by a local superposition and given the B factors of the dark model, structure factors were calculated with *GEMMI* and scaled to the reference amplitudes in resolution shells, and uncertainties were estimated from maps with randomized signs of the amplitude differences. For 2025, the same simulation overestimates the refined difference in γ-phosphate occupancy between the interleaved and laser-off darks (0.25, Table 2) 1.4-fold, and 2025 values are given for both calibrations. Detection limits were obtained in the same way, from the difference density calculated for complete loss of the γ-phosphate or complete disorder of Jα (residues 522–543) on the reflections, scale and phases of each observed map; the 3σ limit, increased by any observed change in the direction of loss, was divided by the activated fraction. For Jα, the noise of the mean main-chain value was taken from all 22-residue windows of Rac1 in the same map (Table S3).

### Model building and refinement

The 2025 laser-off dark structure was determined by molecular replacement in Phaser (McCoy *et al*., 2007) using the PA-Rac1 structure of Wu *et al*. (2009) (PDB 2WKP) as the search model, and refined to 2.43 Å through iterative cycles of reciprocal-space refinement in PHENIX (Liebschner *et al*., 2019) and manual rebuilding in Coot (Emsley *et al*., 2010), including optimization of side-chain conformations and placement of water molecules. The 2026 interleaved dark structure was determined and refined to 2.01 Å in the same way, also starting from 2WKP. In this model, the γ-phosphate of GTP (PG, O1G, O2G and O3G) was refined as one occupancy group (0.78), consistent with the GDP-bound sub-population of the 2026 interleaved dark (Table 2). These two structures are referred to below as the 2025 and 2026 dark models.

All light-state and GDP-state models were refined against qFextr with phenix.refine (coordinates, isotropic B factors and occupancies). In the light-state models of both experiments only LOV2, FMN, residue 648, the interface Ca²⁺ ion and nearby water molecules were refined, with Rac1, GTP and Mg²⁺ kept at the coordinates of the starting model and the γ-phosphate of GTP at the occupancy of the interleaved dark reference refined as one group in Phenix (0.68 in 2025, 0.78 in 2026; Table 2), Cys450 as a single conformer covalently bonded to FMN C4a, and torsion restraints to the starting model outside the flavin pocket; the 2025 light-state model started from the 2025 dark model, and the 2026 models from per-window models refined earlier in this study against the same data. In every 2026 window, Ile445 was rebuilt in the light-state rotamer (χ1 −65°), which side-chain omit maps supported (+4.3 to +5.0σ, against +1.4 to +3.1σ for the dark-state rotamer). In the 2025 light-state model, Ile445 was rebuilt as a single conformer in the same rotamer (χ1 −65°, χ2 170°), taken from the 2026 0–350 µs model after superposition on LOV2, with its side-chain torsions restrained to that rotamer as in the 2026 models; a polder omit map supports it (+4.4 to +5.1σ at Cγ1, Cγ2 and Cδ1, against +1.8 to +2.6σ at Cγ2 and Cδ1 of the dark-state rotamers). The GDP-state model combined the LOV2 domain of the 2025 dark model with Rac1, GDP and the Mg²⁺ site of an earlier refinement of the same population in this study (event 6), with Mg²⁺–ligand distances restrained to 2.08 Å. The switch I conformation of the GDP-state model was tested with polder omit maps (Liebschner et al., 2017) calculated from the qFextr amplitudes of the GDP pair with residues 569–583 (Rac1 26–40) omitted and, as a reference, with a Rac1 core segment of the same length (residues 702–716) omitted; the maps were calculated with the polder program of cctbx 2026.9 and agreed within 0.1σ with Phenix 1.19.2. To test alternative switch I conformations, the density calculated for the modeled switch I was subtracted from the polder map, and the mean residual over the atoms that distinguish each alternative was compared with that of 1,500 random rigid placements of the same atoms. For switch I of free GDP-bound Rac1 (8S1N and 5N6O, placed by the core superposition), the residual was no higher than for random placements (p = 0.26–0.41), and the mFextr–DFc map had no peak above 2.8σ within 0.8 Å of its Cα atoms. Cys648 (Rac1 Cys105) carries additional density in all datasets and was modeled as S-sulfocysteine in all models (group occupancy 0.49 and 0.62 in the 2025 and 2026 dark models and 0.51–0.70 in the light-state and GDP-state models); its chemical identity is tentative, because the crystallization buffer contains cacodylate. For the 1.00–1.35 ms window, whose original R-free set was not retained, the free set of the 1.50–1.85 ms model was used after a debiasing step, and Rac1, GTP and Mg²⁺ were taken from the 1.50–1.85 ms model (Supplemental Methods). Because only LOV2 was refined against amplitudes extrapolated 6.7- to 12.5-fold, the relative orientation of the two domains in the light-state models is not interpreted; it was determined from the maps (see Rigid-body analysis of the difference maps).

In all structures, electron density could be interpreted for residues 404–723. All deposited models were finalized with phenix.refine 1.21.2 and validated with MolProbity (Williams *et al*., 2018) and the wwPDB validation reports (Table 1).

Coordinates and structure factors have been deposited in the Protein Data Bank under accession codes 35EU, 35EV, 35FB, 35FI, 35FN, 35FK, 35FL and 35FT (Table 1). Each structure-factor file of an extrapolated dataset contains the extrapolated amplitudes used for refinement, with their R-free flags, followed by the merged intensities of the triggered and reference datasets.

### Structure comparison

Structures were superposed on Cα atoms with *GEMMI* (Wojdyr, 2022). For Rac1, the core (fusion residues 547–723 without switch I, switch II and one flanking residue on each side, 568–589 and 599–620) was used; for LOV2, residues 404–521. Estimated maximum-likelihood coordinate errors are 0.25 Å for the 2026 dark model, 0.37 Å for the 2025 dark model, 0.42–0.54 Å for the 2026 light-state models, 0.55 Å for the 2025 light-state model and 0.42 Å for the GDP-state model; differences smaller than these values are not interpreted individually. Molecular figures were prepared with PyMOL (Schrödinger, LLC).

Secondary-structure elements were assigned with DSSP (Kabsch & Sander, 1983) on the 2026 dark model. The PAS core of LOV2 comprises the Aβ (Asn415–Thr418) and Bβ (Ile427–Ala430) strands and the Cα (Asp432–Thr438), Dα (Arg442–Ile444) and Eα (Arg451–Leu453) helices, followed by the helical connector Fα (Arg460–Asp471) and the β-scaffold strands Gβ (Val476–Tyr483), Hβ (Lys489–Arg500) and Iβ (Val506–Gly516). The short Eα element adopts 3₁_0_ geometry and is preceded directly by Cys450. The N-terminal cap (Leu404–Asn414) contains a single 3₁_0_ turn, A′α (Leu407–Arg410), and the C-terminal flanking region comprises the Iβ–Jα loop (Thr517–Arg521) and the Jα helix (Asp522–Ala543). Rac1 has a central six-stranded β-sheet of five parallel and one antiparallel strand surrounded by six α-helices and short 3₁_0_ segments.

### Rigid-body analysis of the difference maps

The relative motion of LOV2 and Rac1 was determined directly from the difference maps, by least-squares fitting of each observed map with maps calculated for small rigid-body rotations and translations of two bodies, LOV2 (residues 404–521 with FMN) and Rac1 (residues 547–723 with GTP and Mg²⁺), with Jα (522–546) excluded. The fit used the observed amplitude differences between 2.5 and 8 Å, the phases and bulk-solvent model of the dark structure, and nonlinear refinement of the fitted motion. The tilt is the rotation of LOV2 relative to Rac1, and its axis is the screw axis of the relative motion; interface atoms are the atoms of either domain within 4 Å of the other. Uncertainties were estimated by bootstrapping reflections, and significance from null maps with randomized signs of the amplitude differences and with noise matched to the fit residuals. The same fit applied to the difference between the two 2026 dark datasets (interleaved minus laser-off) served as a control, and values per activated molecule were obtained by dividing by the activated fraction (see Extrapolation, occupancy and activated fraction). The 2025 map was fitted in the same way, with the phases, bulk-solvent model and fitted model of the 2025 laser-off dark structure. For the light state of the same crystal form accumulated by repeated excitation at an X-ray free-electron laser (XFEL light state; Pachota *et al*., 2026; PDB entries 35HK and 35HL), the fit was made in the same way on the XFEL difference map with the dark model of the XFEL data; on the MicroMAX maps, the remaining differences in amplitude scaling and phases change the fitted tilt by about 2% or less.

### Peak statistics and reproducibility

Local extrema of the difference maps (3 × 3 × 3 grid filter) were assigned to the nearest protein atom within 3 Å. Reproducibility of the 2025 light response was assessed at the 2025 LOV2 extrema of at least 3.5σ, which were scored as recurring when the four 2026 maps, after superposition on LOV2 Cα atoms, had the same sign at 2.5σ or more at that position in at least two of them.

### Comparison with deposited Rac1 structures

Coordinates of GDP-bound Rac1 (PDB entries 8S1N, 5N6O, 1HH4, 1I4D, 2H7V and 2P2L; eleven Rac1 chains) and of GppNHp-bound wild-type Rac1 (3TH5, chain A; chain B lacks part of switch I) were taken from the Protein Data Bank and compared with the fusion models in Rac1 numbering (fusion residue number minus 543). Candidate reporters were taken from a screen of 2,524 interatomic distances and angles across the fusion models and the deposited structures; among the switch-to-core Cα distances, the distance between Ala13 and Tyr32 separated the four time-resolved models most clearly from the deposited GDP-bound structures, and the distances from Thr35 Cβ to the Ala59 carbonyl carbon and from Thr58 Cγ2 to the Thr75 carbonyl carbon were also evaluated; the latter, a switch II reporter, is 11.81–11.90 Å in all fusion models, within the deposited range of 11.43–12.27 Å. Per-residue Cα displacements were calculated after least-squares superposition of the Cα atoms of the Rac1 core (Rac1 residues 4–180 excluding 25–46 and 56–77; 127–133 atoms) onto the 2025 dark model. Electron density at superposed atomic positions was sampled with GEMMI (Wojdyr, 2022) in maps normalized to the r.m.s. of the unit cell. All values were computed from the deposited coordinates and the final fusion models.

### Molecular dynamics simulations

Simulations were started from the 0–350 µs model refined earlier against the 2026 data (its Rac1, GTP and Mg²⁺ coordinates are those of the deposited 0–350 µs entry), which provided the protein, the interface Ca²⁺ ion and the covalent cysteinyl–flavin adduct; the simulated fusion therefore carries the light-state flavin chemistry, whereas the GDP-bound sub-population was observed in crystals without illumination. GTP was replaced by GDP, crystal water molecules were retained, one water molecule was placed in the direction of the vacated γ-phosphate, and no inorganic phosphate was included. The fusion was modeled in explicit CHARMM TIP3P water (Jorgensen et al., 1983) with the CHARMM36m protein force field (Huang *et al*., 2017), CHARMM nucleotide and diphosphate parameters (Pavelites et al., 1997; Foloppe & MacKerell, 2000; Denning et al., 2011), CHARMM ion parameters (Beglov & Roux, 1994) with pair-specific corrections for Ca²⁺–carboxylate and Ca²⁺–chloride/phosphate interactions (Liao et al., 2016; Han et al., 2018) and published parameters for the cysteinyl–flavin adduct (Freddolino *et al*., 2013), and simulated with OpenMM 8.0 (Eastman *et al*., 2024) under periodic boundary conditions, with particle-mesh Ewald electrostatics, a 1.2 nm non-bonded cutoff with switching from 1.0 nm, constrained bonds to hydrogen, a 2 fs time step, Langevin-middle integration at 303.15 K (friction coefficient 1 ps⁻¹) and a Monte Carlo barostat at 1 bar.

After initial biasing (Wei *et al*., 2024), the Mg²⁺ ion adopted a GDP-like coordination, with one GDP oxygen, Thr560 OG1 and four water molecules, which remained stable over 20 ns of unrestrained simulation. Collective variables (interatomic distances and angles, many of them involving side-chain atoms; Data S1) were then selected by the separation of the starting and target ensembles relative to their fluctuations and biased toward the values of an earlier model of the GDP-bound sub-population, refined at 2.60 Å from class 6 of the 2025 illuminated frames, which agrees with the GDP-state model reported here to 0.44 Å (switch I) and 0.15 Å (switch II) Cα after superposition on the Rac1 core; values that had been reached were retained while remaining mismatches were biased further. The final restraint sets comprised 12 distances and seven angles in replica 1 and 32 distances and three angles in replica 2 (Figure S5). The restraints were released linearly over 10 ns (replica 1) or 5 ns (replica 2), and each replica was then followed for 40 ns of unbiased dynamics. Cα RMSDs of switch I (Rac1 residues 28–40) and switch II (Rac1 residues 60–76) from that model were calculated after superposition on Rac1 residues 4–180 excluding both switches; in the final 5 ns of replica 2, Rac1 as a whole deviated more from that model than the switches did (median Cα RMSD 1.16 Å, with 57% of frames below the 95th percentile of the reference simulation). Reference ranges were taken from the final 10 ns of a simulation started from that model, with the same photoadduct parameters.

## Supplemental Methods. R-free set of the 1.00–1.35 ms window

The R-free flags of the per-window refinement of the 1.00–1.35 ms data were not retained. The free set of the 1.50–1.85 ms model was used instead, and the starting model combined LOV2, FMN, the Ca²⁺ ion and nearby water molecules of the 1.00–1.35 ms model with Rac1, GTP, Mg²⁺ and the remaining water molecules of the 1.50–1.85 ms model, which had been refined with this free set. To remove memory of the original free set, the B factors of the atoms taken from the 1.00–1.35 ms model were reset to their mean (50.2 Å²) and their coordinates were randomly displaced by 0.25 Å r.m.s., followed by five macro-cycles of refinement with torsion restraints to the undisplaced model. R_free_ was 0.3329 after this procedure and 0.3315 in a control refinement without it, and apart from Ile445 no side chain changed rotamer.

---

**Figure S1.**
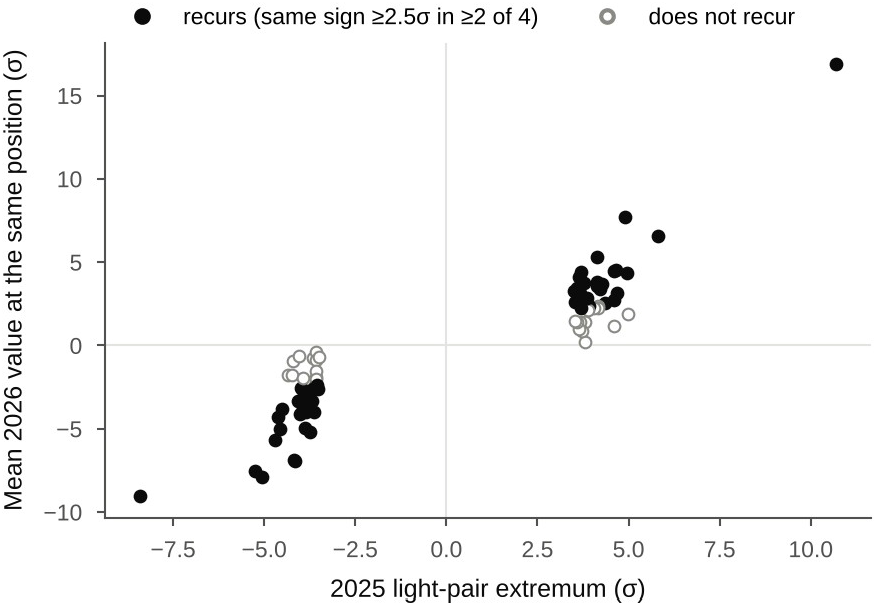
Reproducibility of the 2025 light response. Mean value in the four 2026 maps at the positions of the LOV2 extrema of at least 3.5σ in the 2025 illuminated minus interleaved dark map (light pair), plotted against their 2025 value, after superposition on LOV2; filled circles, extrema that recur with the same sign at 2.5σ or more in at least two of the four 2026 maps.

**Figure S2.**
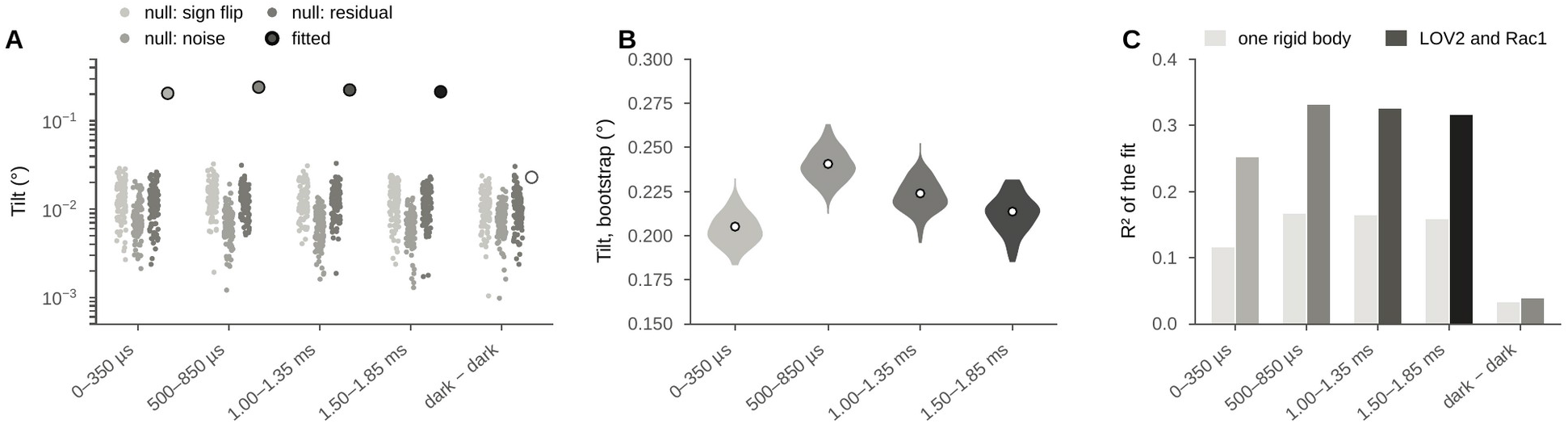
Significance of the rigid-body fits. (A) Fitted tilt (observed, not scaled by the activated fraction) for each 2026 window and the 2026 dark–dark control (circles), with null distributions from maps with sign-flipped amplitude differences, noise maps and residual maps (dots; logarithmic scale). (B) Bootstrap distributions of the tilt for the 2026 windows; circles, fitted values. (C) Fraction of the difference density explained (R²) with LOV2 and Rac1 fitted as one rigid body (light bars) or as two bodies (dark bars).

**Figure S3.**
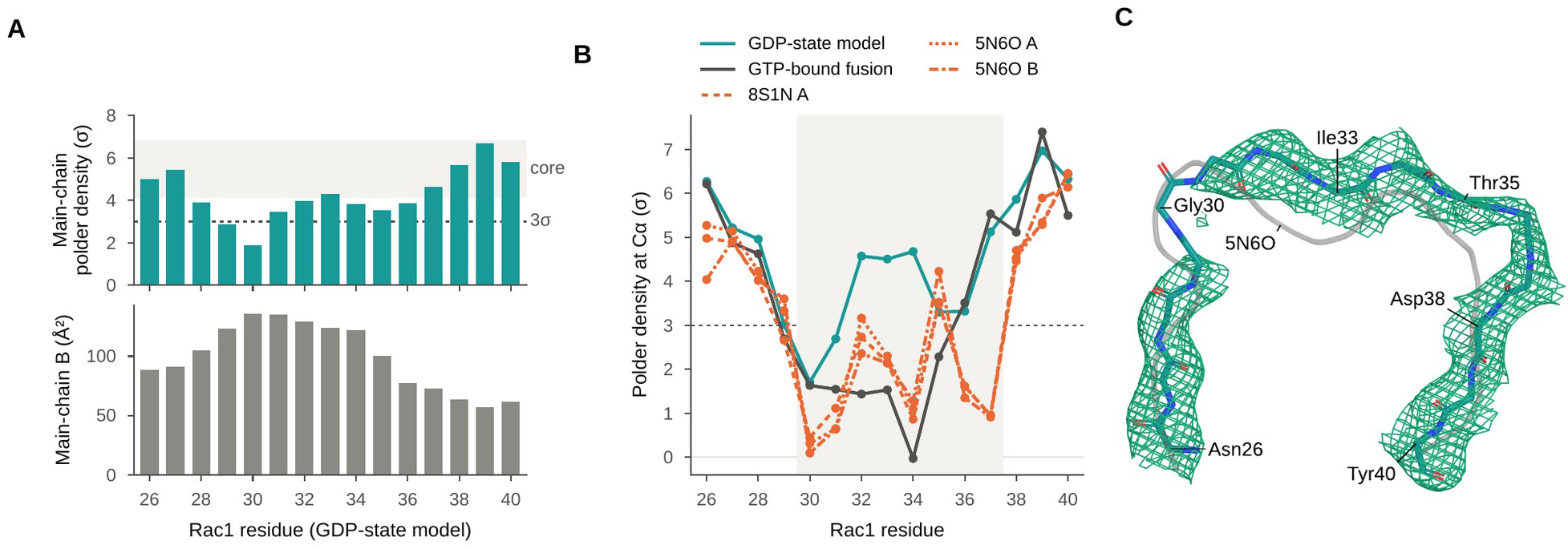
Switch I in the GDP-state model. (A) Mean main-chain density per residue in a polder omit map with residues 569–583 (Rac1 26–40) omitted, and mean main-chain B factor; shaded band, range for a Rac1 core segment of the same length (residues 702–716, Rac1 159–173) omitted in the same way; dashed line, 3σ. (B) Polder density at the Cα positions of the GDP-state model, of the GTP-bound fusion (2025 laser-off dark model, as refined in the same crystal frame) and of free GDP-bound Rac1 (8S1N chain A, 5N6O chains A and B) placed by superposition on the Rac1 core; shaded, Gly30–Phe37; dashed line, 3σ. (C) Polder map at 3.0σ (green), carved 2.0 Å around the switch I main chain of the GDP-state model (teal sticks), with switch I of 5N6O chain A (gray tube); labels in Rac1 numbering. The maps were calculated from the qFextr amplitudes of the GDP pair (occupancy 0.25, 2.6 Å; Methods); σ is the r.m.s. of the whole unit cell of each map.

**Figure S4.**
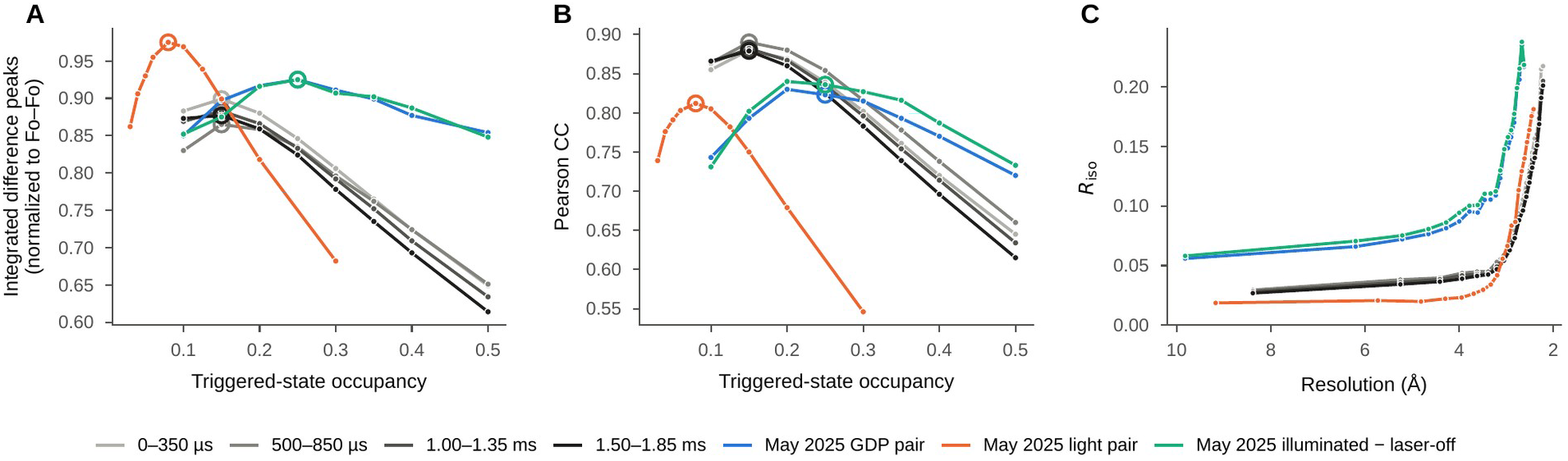
Occupancy determination in Xtrapol8 and isomorphism for all seven difference pairs. (A) Normalized integrated difference-map signal and (B) Pearson correlation between extrapolated and Fo–Fo maps against the assumed triggered-state occupancy; open circles mark the selected values (0.15 for each 2026 window; 0.08, 0.25 and 0.25 for the three 2025 pairs). (C) R_iso_ by resolution shell.

**Figure S5.**
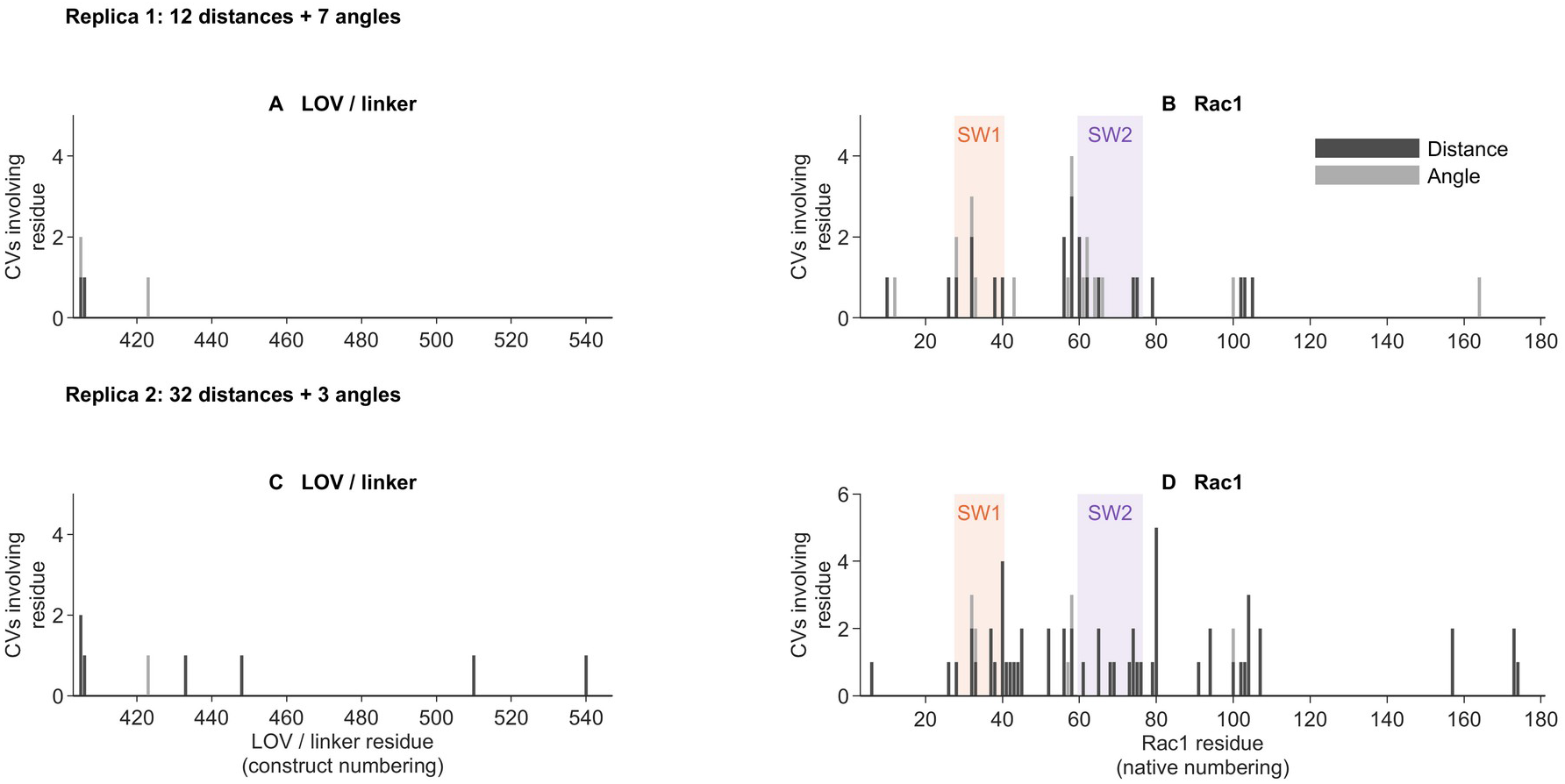
Residues in the restraint sets retained in the two molecular dynamics simulations. Each distance (dark gray) or angle (light gray) restraint contributes one count to each participating residue. Replica 1 (A, B) used 12 distances and seven angles, replica 2 (C, D) 32 distances and three angles. LOV2 and linker residues are numbered as in the fusion, Rac1 residues as in Rac1; switch I (Rac1 28–40) and switch II (Rac1 60–76) are shaded orange and purple, respectively. Counts describe the applied restraints, not the importance of a residue. Atom definitions are given in Data S1.

**Table S1.**
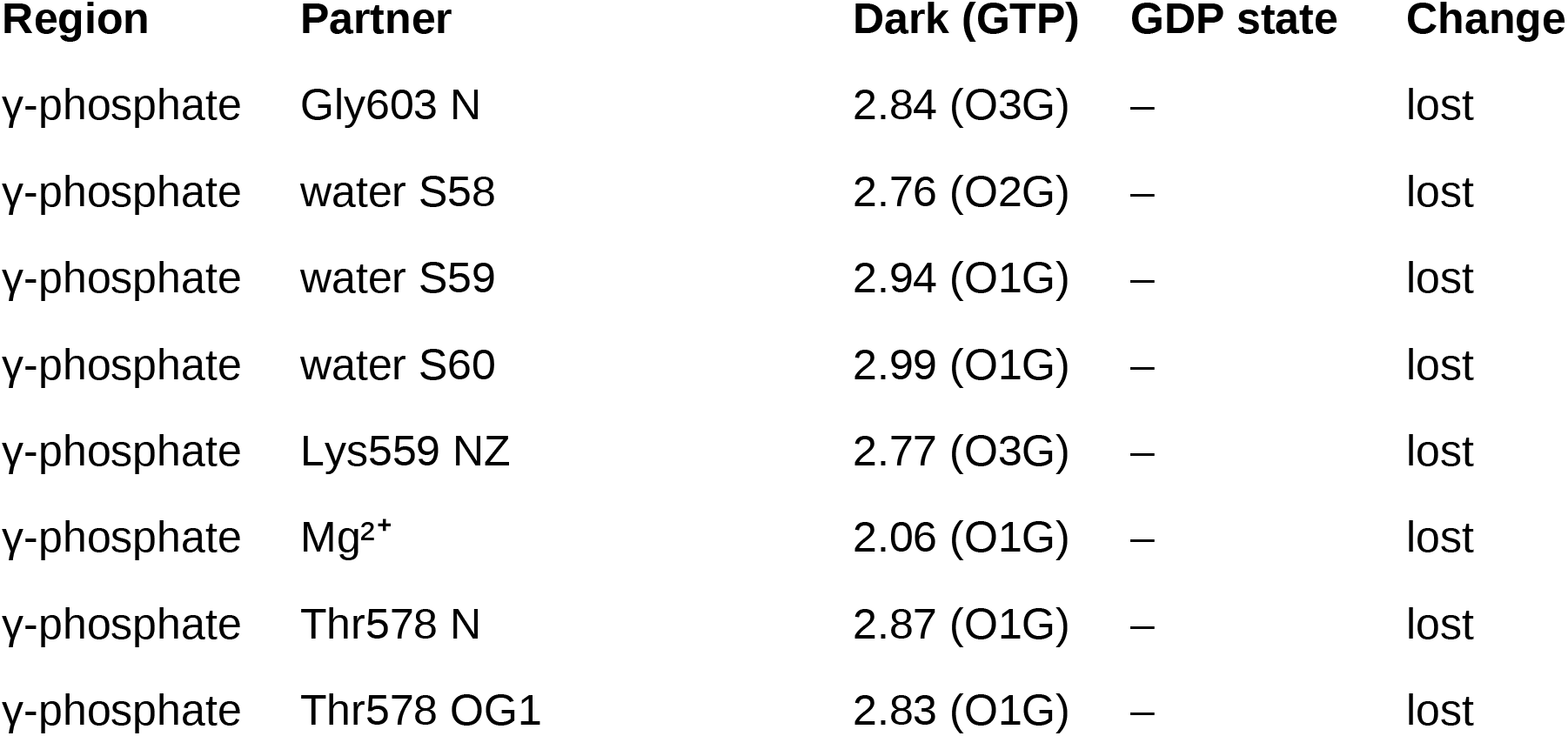
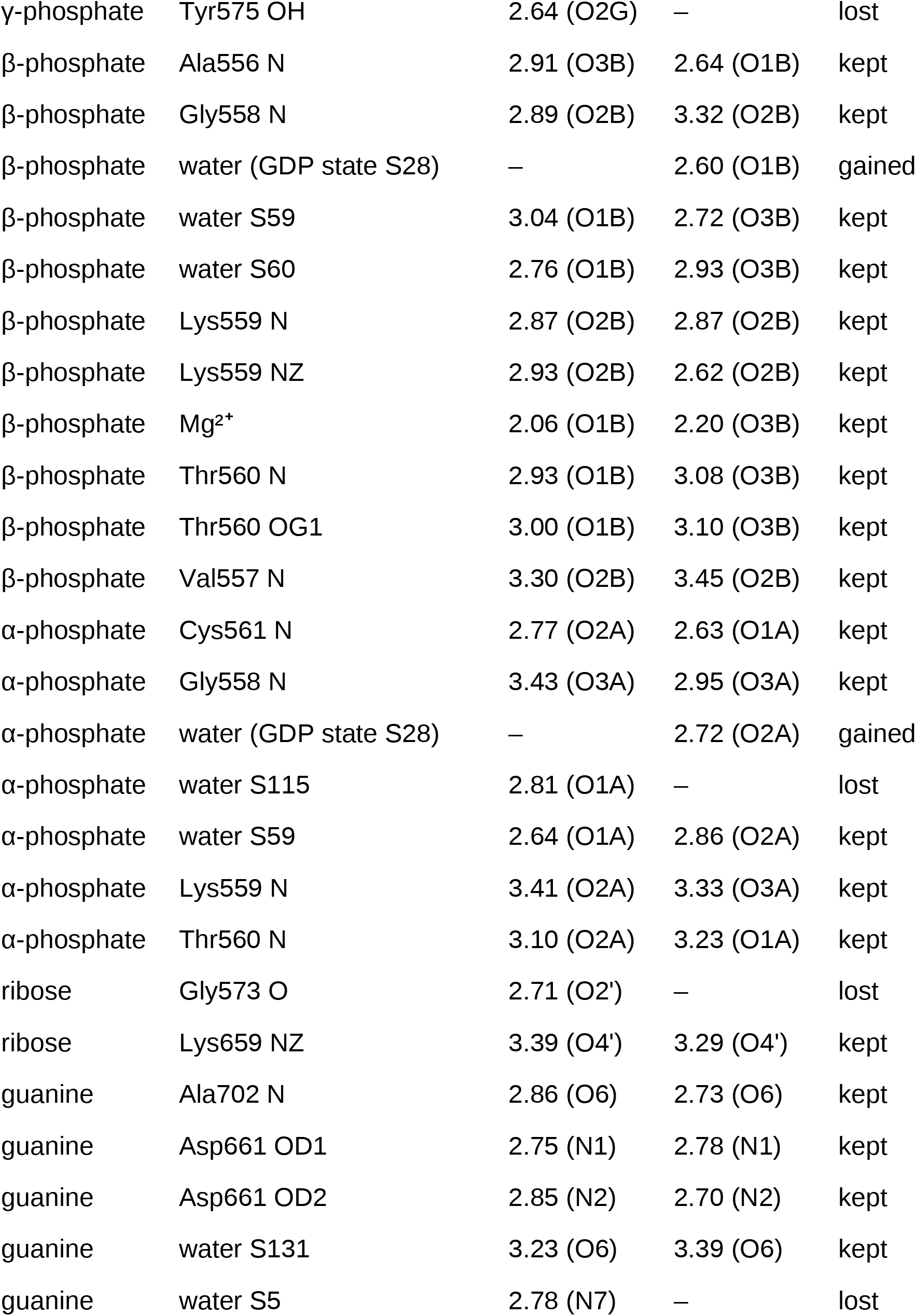
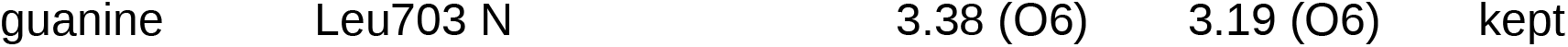
Polar contacts (≤ 3.5 Å) between the nucleotide and its environment in the 2025 dark (GTP) and GDP-state models (shortest contact per partner atom and region, Å).

**Table S2.**
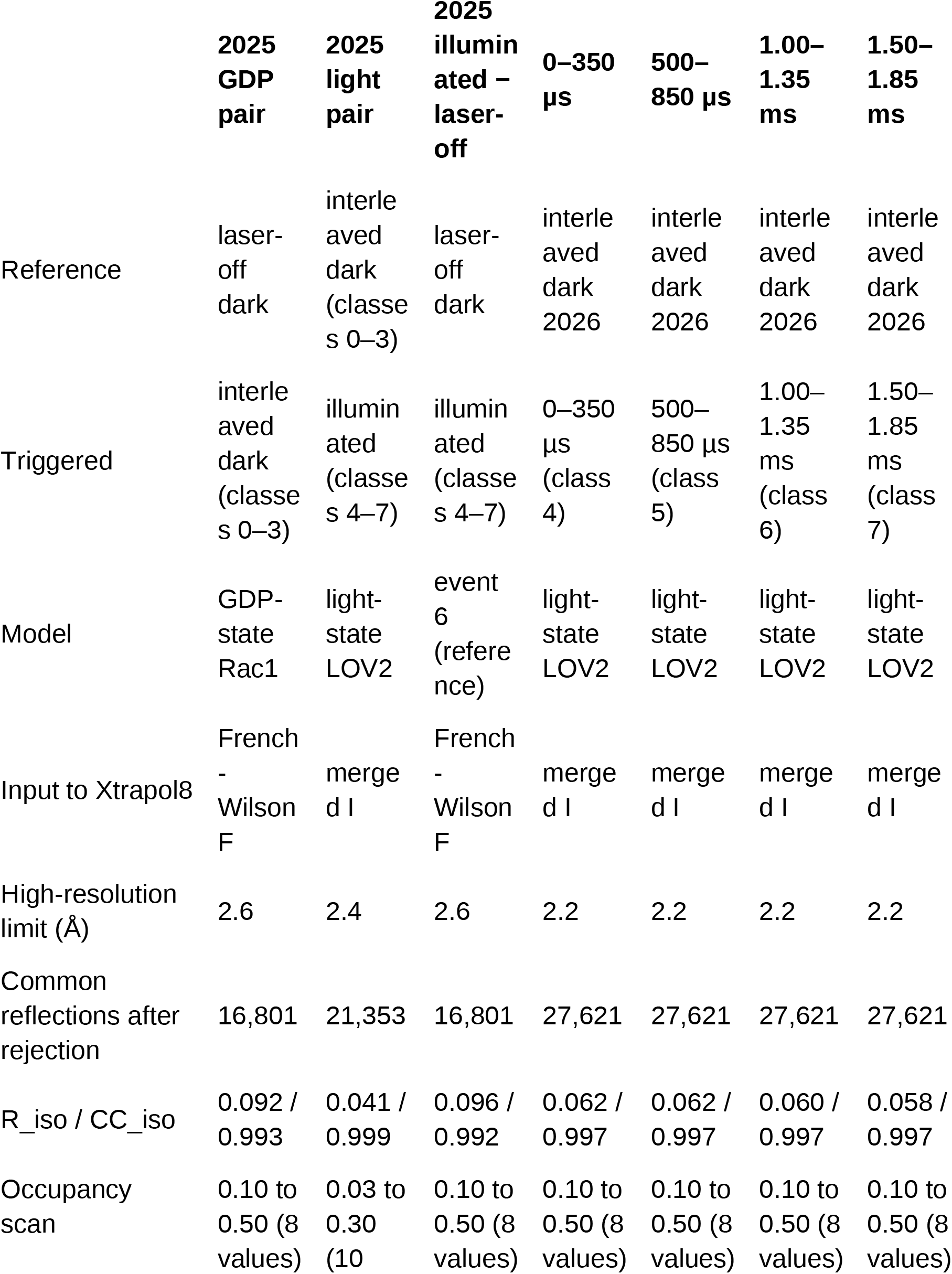
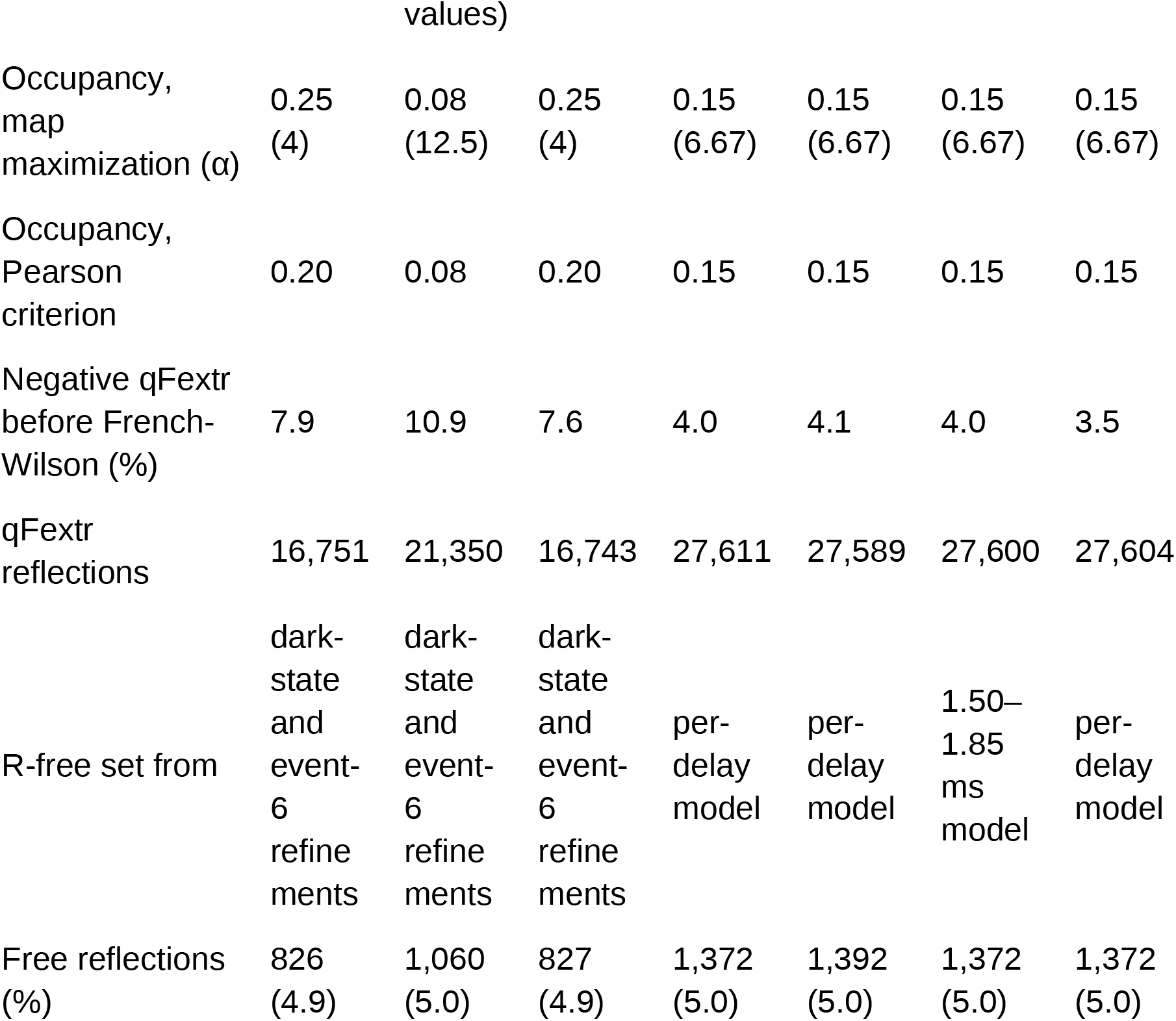
Xtrapol8 extrapolation for each difference pair.

**Table S3.** Activated fraction and detection limits for each difference map (unweighted maps phased with the dark model, 2.2–8 Å for 2026 and 2.4–8 Å for 2025; Methods). Activated fraction with the standard deviation from maps with randomized signs of the amplitude differences, and correlation between observed and simulated density over the chromophore pocket. Bounds are 3σ limits per activated molecule, increased by any observed change in the direction of the effect. For 2025, values in parentheses use the empirical calibration from the GDP pair (Methods).

| Map | Activate<br>d<br>fraction | CC in<br>pocke<br>t | γ-<br>Phosph<br>ate loss<br>detectab<br>le at $3\sigma$<br>(% of<br>molecul<br>es) | Observe<br>d γ-<br>phospho<br>rus ($\sigma$) /<br>implied<br>loss (%<br>of<br>molecule | Bound<br>on γ-<br>phosph<br>ate loss<br>(% of<br>activate<br>d<br>molecul | Bound<br>on $J\alpha$<br>disorder<br>(% of<br>activate<br>d<br>molecul<br>es) |
| --- | --- | --- | --- | --- | --- | --- |

|  |  |  |  | s) | es) |  |
| --- | --- | --- | --- | --- | --- | --- |
| 0–350 μs –<br>interleaved<br>dark | $0.20 \pm$<br>0.01 | 0.82 | 3.2 | –3.3 /<br>+3.5 | 34 | 17 |
| 500–850 μs –<br>interleaved<br>dark | $0.24 \pm$<br>0.01 | 0.85 | 3.2 | –1.0 /<br>+1.1 | 18 | 11 |
| 1.00–1.35 ms<br>– interleaved<br>dark | $0.23 \pm$<br>0.01 | 0.85 | 3.1 | –0.8 /<br>+0.8 | 17 | 14 |
| 1.50–1.85 ms<br>– interleaved<br>dark | $0.22 \pm$<br>0.01 | 0.85 | 3.0 | +0.3 /<br>–0.3 | 14 | 13 |
| 2025<br>illuminated –<br>interleaved<br>dark | $0.09 \pm$<br>0.01<br>(0.07) | 0.73 | 2.0 (2.8) | +1.4 /<br>–0.9 | 22 (42) | 18 (34) |
| 2026<br>interleaved<br>dark – laser-off<br>dark (control) | $-0.006 \pm$<br>0.006 | –0.07 | – | – | – | – |

**Table S4.**
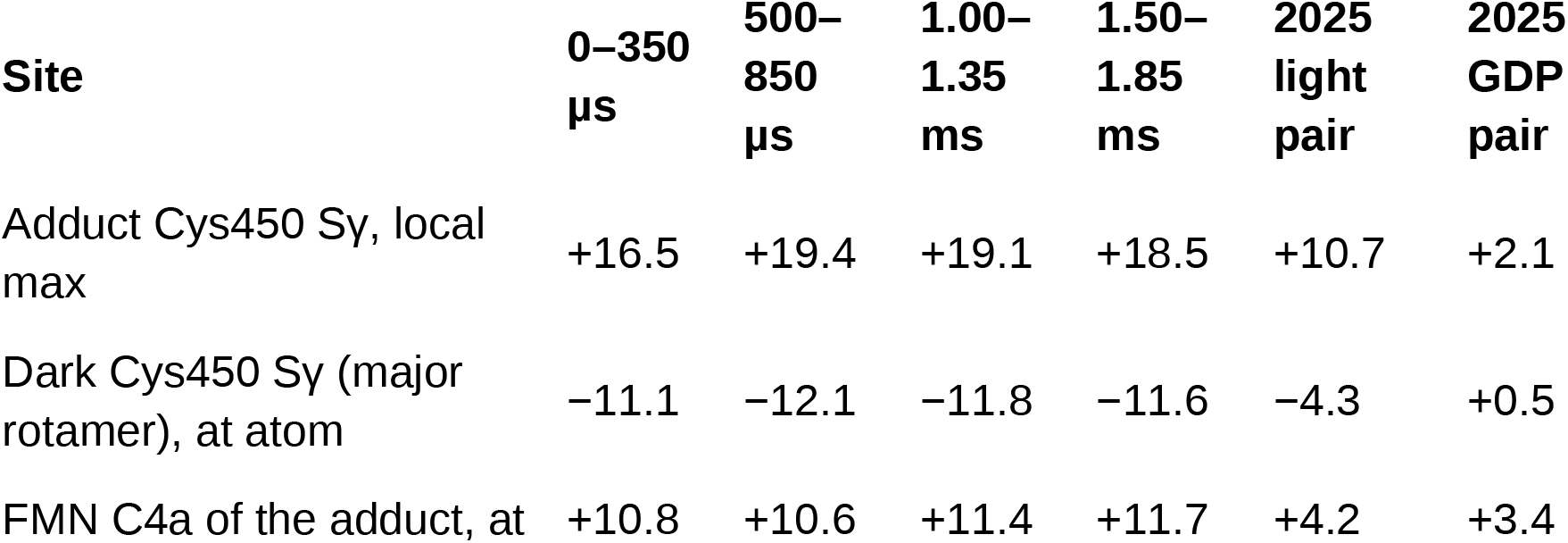
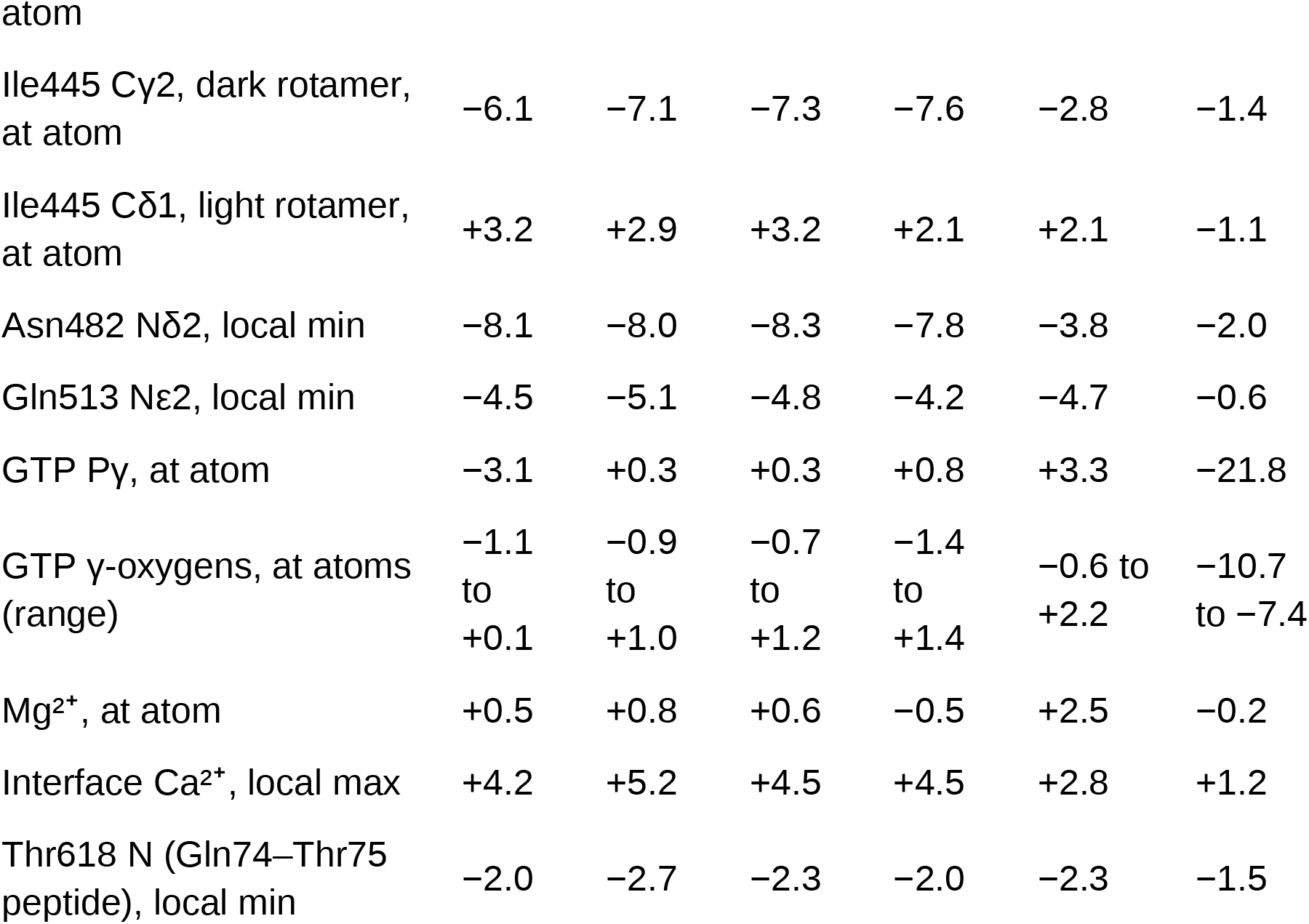
Difference-map values at selected sites (σ, whole unit cell; Xtrapol8 maps, sample rate 4, tricubic interpolation). 2026: each window minus the interleaved dark; 2025 light pair: illuminated minus interleaved dark; 2025 GDP pair: interleaved dark minus laser-off dark. Positions of the adduct and its flavin in the 2025 maps are taken from the 2026 0–350 µs model after a local superposition on residues 444–456, and the Ile445 light rotamer after superposition on LOV2 (404–521).

| Site | 0–350<br>μs | 500–<br>850<br>μs | 1.00–<br>1.35<br>ms | 1.50–<br>1.85<br>ms | 2025<br>light<br>pair | 2025<br>GDP<br>pair |
| --- | --- | --- | --- | --- | --- | --- |
| Adduct Cys450 Sy, local<br>max | +16.5 | +19.4 | +19.1 | +18.5 | +10.7 | +2.1 |
| Dark Cys450 Sy (major<br>rotamer), at atom | –11.1 | –12.1 | –11.8 | –11.6 | –4.3 | +0.5 |
| FMN C4a of the adduct, at | +10.8 | +10.6 | +11.4 | +11.7 | +4.2 | +3.4 |
| Ile445 Cy2, dark rotamer,<br>at atom | -6.1 | -7.1 | -7.3 | -7.6 | -2.8 | -1.4 |
| Ile445 Cδ1, light rotamer,<br>at atom | +3.2 | +2.9 | +3.2 | +2.1 | +2.1 | -1.1 |
| Asn482 Nδ2, local min | -8.1 | -8.0 | -8.3 | -7.8 | -3.8 | -2.0 |
| Gln513 Nε2, local min | -4.5 | -5.1 | -4.8 | -4.2 | -4.7 | -0.6 |
| GTP Py, at atom | -3.1 | +0.3 | +0.3 | +0.8 | +3.3 | -21.8 |
| GTP γ-oxygens, at atoms<br>(range) | -1.1<br>to<br>+0.1 | -0.9<br>to<br>+1.0 | -0.7<br>to<br>+1.2 | -1.4<br>to<br>+1.4 | -0.6 to<br>+2.2 | -10.7<br>to -7.4 |
| Mg<sup>2+</sup>, at atom | +0.5 | +0.8 | +0.6 | -0.5 | +2.5 | -0.2 |
| Interface Ca<sup>2+</sup>, local max | +4.2 | +5.2 | +4.5 | +4.5 | +2.8 | +1.2 |
| Thr618 N (Gln74–Thr75<br>peptide), local min | -2.0 | -2.7 | -2.3 | -2.0 | -2.3 | -1.5 |

**Table S5.** Mg²⁺ coordination (≤ 2.6 Å) in the 2025 dark (GTP) and GDP-state models.

| Dark (GTP) ligand | Distance<br>(Å) | GDP-state ligand | Distance<br>(Å) |
| --- | --- | --- | --- |
| GTP O1B (β-phosphate) | 2.06 | GDP O3B (β-phosphate) | 2.20 |
| Thr560 OG1 (Rac1 Thr17) | 2.08 | Thr560 OG1 (Rac1 Thr17) | 2.12 |
| GTP O1G (γ-phosphate) | 2.06 | water | 2.05 |
| Thr578 OG1 (Rac1 Thr35) | 2.08 | water | 2.10 |
| water | 2.08 | water | 2.14 |
| water | 2.13 | water | 2.17 |

